# Selection of single domain anti-transferrin receptor antibodies for blood-brain barrier transcytosis using a neurotensin based assay and histological assessment of target engagement in a mouse model of Alzheimer’s related amyloid-beta pathology

**DOI:** 10.1101/2022.05.02.490319

**Authors:** Shiran Su, Thomas J. Esparza, David L. Brody

## Abstract

The blood-brain barrier (BBB) presents a major obstacle in developing specific diagnostic imaging agents for many neurological disorders. In this study we aimed to generate single domain anti-mouse transferrin receptor antibodies (anti-mTfR VHHs) to mediate BBB transcytosis as components of novel MRI molecular contrast imaging agents. Anti-mTfR VHHs were produced by immunizing a llama with mTfR, generation of a VHH phage display library, immunopanning, and *in vitro* characterization of candidates. Site directed mutagenesis was used to generate additional variants. VHH fusions with neurotensin (NT) allowed rapid, hypothermia-based screening for VHH-mediated BBB transcytosis in wild-type mice. One anti-mTfR VHH variant was fused with an anti-amyloid-beta (Aβ) VHH dimer and labeled with fluorescent dye for direct assessment of *in vivo* target engagement in a mouse model of AD-related Aβ plaque pathology. An anti-mTfR VHH called M1 and variants had binding affinities to mTfR of <1nM to 1.52nM. The affinity of the VHH binding to mTfR correlated with the efficiency of the VHH-NT induced hypothermia effects after intravenous injection of 600 nmol/kg body weight, ranging from undetectable for nonbinding mutants to −6°C for the best mutants. The anti-mTfR VHH variant M1_P96H_ with the strongest hypothermia effect was fused to the anti-Aβ VHH dimer and labeled with Alexa647; the dye-labeled VHH fusion construct still bound both mTfR and Aβ plaques. However, after intravenous injection at 600 nmol/kg body weight into APP/PS1 transgenic mice, there was no detectible labeling of plaques above control levels. Thus, NT-induced hypothermia did not correlate with direct target engagement in cortex. There was a surprising dissociation between NT-induced hypothermia, presumably mediated by hypothalamus, and direct engagement with Aβ-plaques in cortex. Alternative methods to assess anti-mTfR VHH BBB transcytosis will need to be developed for anti-mTfR VHH screening and the development of novel MRI molecular contrast agents.

## Introduction

Alzheimer’s disease (AD) is one of the most important causes of dementia in the elderly [1]. About 6.2 million Americans are living with AD, and it’s predicted that the number will increase to 13.8 million by 2050 [2]. With no effective therapies to cure or inhibit significant AD symptom progression [3], AD severely decreases patients’ quality of life and creates an enormous burden on the health care system and society [1, 4]. Currently, clinical AD diagnosis is based on cognition and the relative impact of impairments on daily activities [5]. However, multiple neurodegenerative and vascular pathologies can coexist and produce cognitive and behavioral symptoms which could overlap with each other [6]. This makes it difficult to accurately identify pathology based solely on clinical symptoms. The accuracy of clinical diagnosis of AD at the National Institute of Aging and National Institute of Aging sponsored AD centers varies depending on the clinical and neuropathologic criteria used [7]. The sensitivity of AD diagnosis ranges from 70.9% to 87.3%, whereas specificity ranges from 44.3% to 70.8%, which need to be improved. AD has a very long prodrome stage before clinically observable symptoms [8]. Early diagnosis is preferred to help with early intervention and the development of preventive therapeutics to slow AD development [5]. Detection of patients in the preclinical stages of AD can also help to provide diagnostic information, monitor disease progression, and even monitor the effect of newly developed treatment methods. Imaging methods including MRI and PET have been developed to aid the diagnosis of AD [9, 10]. However, PET imaging for AD provides limited resolution and requires radiation exposure. Structural MRI lacks specificity and does not allow direct visualization of Aβ or tau, the histological hallmarks of AD. There is an unmet need for developing methods like molecular contrast MRI which have better resolution than PET and better specificity than structural MRI imaging.

The blood-brain barrier (BBB) represents a significant obstacle in delivering diagnostic and therapeutic agents to the central nervous system (CNS), preventing uptake of more than 98% of potential neurotherapeutics to brain [11–14]. The BBB consists of endothelial cells held together by tight junctions which hinder paracellular passage. Most molecules do not transfer from blood to brain through the BBB, which protects the brain from toxicity and maintains brain homeostasis. Several methods have been developed to improve the transport of diagnostic and therapeutic agents into the CNS. For instance, the BBB may be temporarily opened by administration of hypertonic agents or focused ultrasound. Alternatively, very high doses of an agent can be given so that even if a small fraction enters the brain the desired effect will be achieved. In some cases, direct injection of agents into the cerebrospinal fluid can be employed [14]. However, these methods are invasive and have a risk of causing infection, toxicity, and neurological dysfunction [13, 14].

BBB crossing based on receptor mediated transcytosis (RMT) is potentially especially promising. The use of protein shuttles has the potential to facilitate the transport of therapeutic agents across the BBB using specific endogenous receptor systems. Candidate receptor systems including transferrin receptors (TfR), low-density lipoprotein receptors, insulin receptors and neuropeptide receptors are highly expressed on the BBB where they mediate receptor-mediated transcytosis [13]. Among the different receptors, TfR has been widely used for transporting macromolecules across the BBB [15, 16]. Based on the study of Yu et al., there is a nonlinear relationship between an antibody’s affinity for TfR and its uptake in brain. At tracer doses, antibodies with higher affinity to TfR have higher uptake into the brain, while at therapeutic doses, antibodies with lower affinity to brain have higher uptake into the brain [17]. The effect of TfR affinity on brain uptake has been confirmed by the study of Wiley et al [18]. Transferrin conjugated gold nanoparticles with high avidity to TfR remain strongly attached to brain endothelial cells and reduced accumulation in brain parenchyma compared with nanoparticles with lower avidity to TfR [18]. Thus, to achieve optimal brain uptake, it is important to optimize anti-TfR concentration and antibody affinity to TfR. Jefferies et al. identified a monoclonal antibody OX-26 specific for transferrin receptors [19]. This antibody was tested and was confirmed to be able to facilitate TfR-mediated transcytosis across BBB [20]. Kissel et al. produced monoclonal antibody 8D3 which recognizes murine transferrin receptor [21]. Yu et al. generated a bispecific antibody that binds to TfR for transcytosis and also to the enzyme β-secretase for inhibiting Aβ production[17]. Hultqvist et al. attached a single chain variable fragment against TfR to the anti-amyloid-beta protofibril recombinant monoclonal antibody RmAb158. The anti-TfR single chain variable fragment increased brain uptake of the antibody by 80-fold [22]. However, most of the existing anti-TfR antibodies have relatively large size, especially when conjugated to additional components for RMT or other payloads such as imaging contrast agents. Previous research found that reducing the size of nanoparticles helped to improve blood brain barrier transcytosis [23] Keeping the size of each component small is important since the ultimate goal of this line of investigation is to develop multicomponent brain MRI molecular contrast agents. Also, it is expensive to engineer and synthesize monoclonal antibodies, which are typically produced in mammalian cell culture. Thus, there is an unmet need for smaller, less expensive, and easier to engineer system for BBB transcytosis.

Camelids produce functional antibodies devoid of light chains called heavy chain-only antibodies (HCAbs) [24, 25]. HCAbs recognize their cognate antigens by one single domain, the variable domain (VHH). The VHH in isolation is very small compared with other antibodies. The molecular weights of VHHs are typically ∼15kDa, which is about 1/10 of the molecular weight of a conventional IgG and about half the molecular weight of a single chain variable fragment (Scfv) [25, 26]. VHHs have affinities at the same order of magnitude as conventional IgGs, often in the nanomolar or subnanomolar range [27]. Because of their small size, VHHs can also bind to epitopes not recognized by conventional antibodies and can have better tissue penetration capacities [28, 29]. The factors governing VHH immunogenicity are similar to those for conventional antibodies and VHHs have been demonstrated to have low immunogenicity risk profile [30, 31]. Caplacizumab was the first VHH approved by FDA for treatment of acquired thrombotic thrombocytopenic purpura in humans [32]. There are several more VHHs which are in clinical trials, with safety profiles similar to other antibody therapeutics in humans [33, 34]. Importantly, VHHs have been found that can facilitate BBB penetration and allow brain target binding. The use of VHHs for BBB transcytosis and target engagement is promising. Specifically, Stanimirovic et al. identified an insulin-like growth factor 1 receptor binding VHH which crosses the BBB by receptor mediated transcytosis [35, 36]. Danis et al. identified and optimized VHHs to mitigate brain accumulation of pathological tau in a tauopathy mouse model [37]. Dupre et al., identified VHHs which could be used to detect tau in transgenic mice brain tissues [38]. However, the extent to which VHHs that engage in receptor mediated transcytosis using the TfR can carry diagnostic and therapeutic payloads across the BBB still has not been assessed. Here we tested the hypothesis that VHHs that bind to TfR and cross the BBB through RMT in mice could be coupled with additional VHHs that bind to amyloid plaques as a proof of concept for a platform which could be generalized to other neurological diseases.

Current methods (ELISA or radioisotope detection) to evaluate BBB crossing require substantial resources and can be time-consuming [17, 39, 40]. An efficient way to screen for BBB crossing *in vivo* would be helpful since *in vitro* models may not be fully predictive of *in vivo* results [41]. Neurotensin (NT) is a 13 amino acid peptide first identified by Carraway and Leeman in 1973 from bovine hypothalamic extracts [42]. NT is expressed in CNS as well as in peripheral tissues, mostly in the gastrointestinal tract. NT is involved in regulating appetite, nociception, and thermoregulation in the CNS, and alters nutrient absorption, gastrointestinal motility, and secretion in the peripheral gastrointestinal tract [43]. It was found that NT induces rapid and transient hypothermia in mice and rats when injected to CNS [44, 45]. The hypothermia was likely to be due to effects in hypothalamus. Young & Kuhar found that NT receptors had moderate to high densities in hypothalamus [46]. Injection of NT to medial, lateral preoptic area and anterior parts of hypothalamus induced the hypothermia effect [46]. Meanwhile, intravenously injected NT does not typically cause hypothermia [47]. On the other hand, when NT-conjugated mouse TfR (mTfR)-binding VHHs were injected intravenously to mice, the NT-mTfR VHH conjugates induced hypothermia in mice, presumably because they get across the BBB and bind to NT receptors in hypothalamus [39, 48]. These properties of the NT system make it an apparently attractive assay platform for rapid testing of VHH BBB transcytosis.

Here, we independently generated additional mTfR-binding VHHs and used a similar NT based modular system to screen these VHHs for mTfR-mediated BBB transcytosis (**Fig 1**). We hypothesized that this modular screening system can be used to identify and optimize anti-mTfR VHHs for BBB transcytosis and brain target binding.

**Fig 1.**
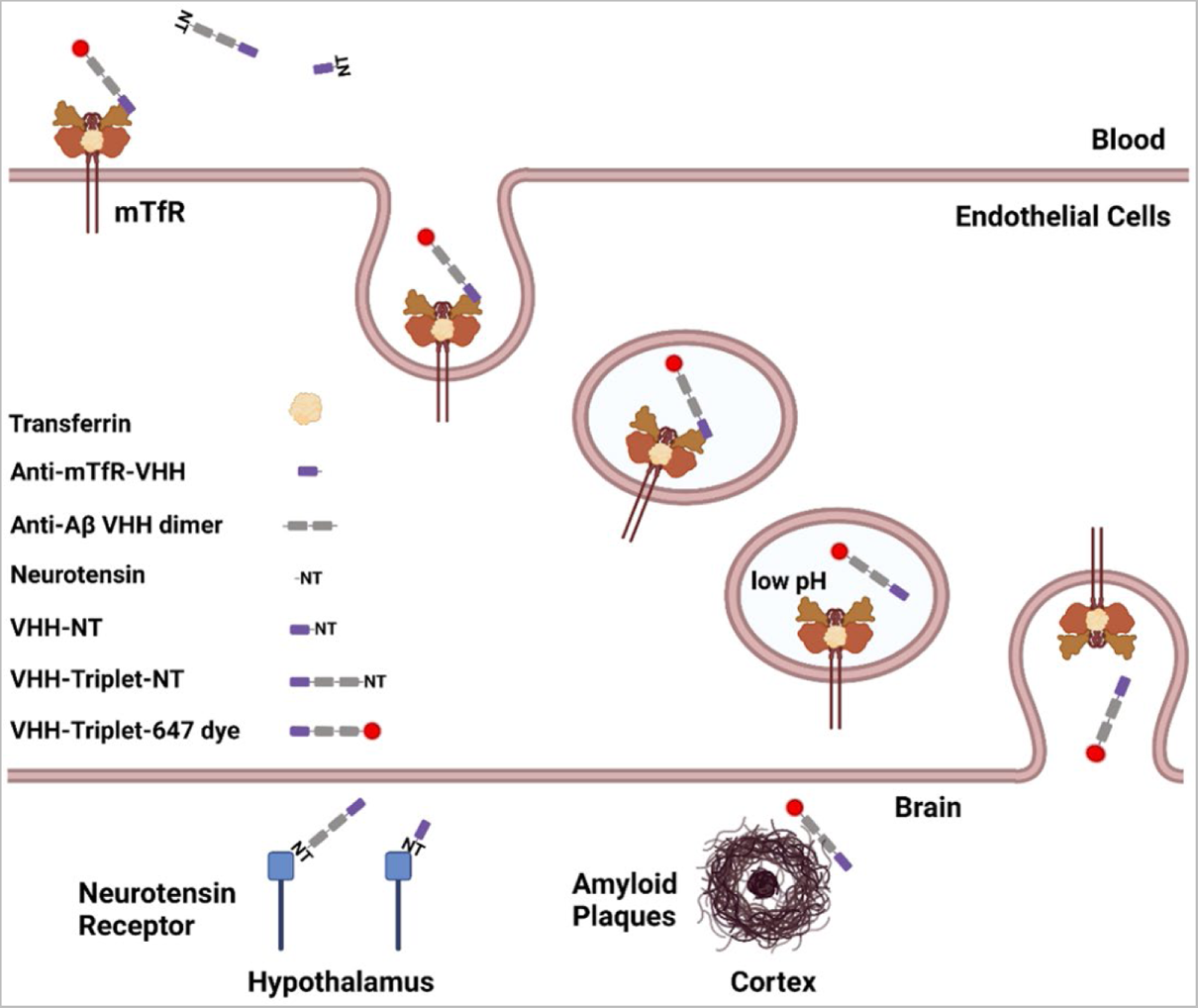
Schematic of the modular system for VHH-mediated BBB transcytosis and target engagement. Module 1: anti-mTfR VHH for RMT across BBB. Module 2: neurotensin peptide (NT) for rapid assessment of *in vivo* target engagement via measurement of hypothermia. Module 3: anti-Aβ VHH dimer for disease-relevant target engagement in brain parenchyma. Module 4: 647 dye for visualization of VHH constructs in situ. The anti-mTfR VHH was combined with NT for anti-mTfR VHH screening through monitoring hypothermia effect. The anti-mTfR VHH were combined with VHH dimer and NT or 647 to test the effect of anti-mTfR VHH together with other components of the module. This schematic figure also shows the TfR mediated transcytosis across BBB. The anti-mTfR VHH conjugates binds to TfR on the endothelial cells which comprise a major portion of the BBB. Then the TfR-VHH complex is endocytosed across the endothelial cells in the endosomes. VHHs dissociate from the TfR-VHH complex with decreased pH level in endosome and are released on the brain side of the BBB. There were two different targets in the brain parenchyma in this study, the NT receptor in hypothalamus and amyloid plaques in cortex and hippocampus. (Figure generated using BioRender.com)

This screening system includes four modules. The first module is an anti-mTfR VHH for receptor mediated transcytosis across the BBB; the second module is the neurotensin peptide for assessment of target engagement *in vivo* through measurement of hypothermia. The third module is a dimer of anti - Aβ VHHs for disease-relevant brain target engagement. The fourth module is a fluorescence dye for visualization of VHH constructs in situ. We tested different VHH variants and found one variant which showed good BBB penetration based on this screening system. The VHH variant with the best BBB transcytosis ability fused to a tandem VHH dimer called Nb3-Nb3 which binds to amyloid plaques in brain parenchyma. This VHH triplet was conjugated to a fluorescent dye and post-mortem confocal microscopy was performed to directly evaluate brain target engagement.

## Materials and Methods

### Immunization of llama with mouse transferrin receptor

A single adult male llama (*Lama glama*) was immunized under contract agreement through Triple J Farms (Kent Laboratories, Bellingham, WA) following the method previously described [49]. Briefly, subcutaneous injections of 100 µg ectodomain (Cys89-Phe763) mTfR (50741-M07H, SinoBiological) (synthesized in HEK293 cells and went through glycosylation) were performed with protein emulsified with complete Freund’s adjuvant on day 0, followed by additional 100 µg immunizations emulsified with incomplete Freund’s adjuvant on days 14, 28, and 42. On day 49, peripheral blood was drawn for peripheral blood mononuclear cell (PBMC) isolation. Triple J Farms operates under established National Institutes of Health Office of Laboratory Animal Welfare Assurance certification number A4335-01 and United States Department of Agriculture registration number 91-R-0054.

### Generation of VHH immune phage display library

The generation of an immune phage display library and isolation of mTfR binding VHH clones was performed using the methods previously described [49]. Briefly, total RNA extracted from PBMCs was used for synthesis of first-strand complimentary DNA (cDNA) using the SuperScript IV First-Strand Synthesis kit (#1891050, Invitrogen). The heavy-chain variable domain was then amplified from the cDNA using Q5 high-fidelity DNA polymerase (New England Biolabs) with the described primers (CALL001: 5′-GTCCTGGCTGCTCTTCTACAAGG-3′ and CALL002: 5′-GGTACGTGCTGTTGAACTGTTCC-3’). The heavy-chain specific amplicon was isolated using electrophoresis with low-melting point agarose extraction with the QIAquick Gel Extraction kit (Qiagen). A secondary amplification was performed using a modification of the primers (VHH-Esp-For: 5′-CCGGCCATGGCTGATGTGCAGCTGCAGGAGTCTGGRGGAGG-3′ and VHH-Esp-Rev: 5′-GTGCGGCCGCTGAGGAGACGGTGACCTGGG T-3′) used by Pardon et al. to facilitate cloning into the phagemid pHEN2 [50]. The amplified sequences were cleaved with the restriction endonucleases NcoI and NotI (New England Biolabs) and subsequently ligated into compatibly cleaved pHEN2 phagemid at a 3:1 (insert:phagemid) ratio overnight at 16°C followed by purification. The resulting ligation mixture was electroporated into TG-1 phage-display competent cells (#60502-1, Lucigen) and plated onto 2xYT agar containing 100 µg/mL carbenicillin and 2% (w/v) glucose at 37°C overnight. The resulting library contained > 10^7^ independent clones. Phage was produced for screening using the M13KO7 helper phage (#18311019, Invitrogen) followed by precipitation by addition of one-fifth volume 20% polyethylene glycol 6000 / 2.5 M sodium chloride solution on ice and centrifugation to purify the phage particles.

### Immunopanning and clone screening

Selection of mTfR specific VHH was performed using direct binding of phage to immobilize mTfR. Standard radioimmunoassay (RIA) tubes were coated with 500 µL mTfR solution at 5 µg/mL in sodium carbonate buffer, pH 9.6 overnight at 4°C. The coating solution was removed, and the RIA tube filled with a 2% (w/v) non-specific blocking solution (bovine serum albumin or nonfat dry milk) in 1x Phosphate Buffered Saline (PBS). Amplified phage (∼10^11^ phage) was mixed with blocking solution to a final volume of 500 µL and then transferred into the RIA tube to allow for association at room temperature and 600 rpm mixing. The RIA tube was then washed 20 times with 1x PBS and then the bound phage eluted with 100 mM triethylamine solution for 20 mins. The eluted phage solution was neutralized with 1:10 volume 1 M Tris-HCl, pH 8.0. The eluted phage was amplified in TG-1 cells and a second round of immunopanning was performed.

Following the second round of immunopanning, individual colonies were selected and cultured in 96-well blocks containing 2xYT containing carbenicillin at 37°C with 300 rpm shaking for 4-6 hours. Expression of VHH was induced by addition of isopropyl-beta-D-thiogalactoside (IPTG) to a final concentration of 1 mM and incubation overnight at 37°C. The culture blocks were centrifuged to pellet the cells and frozen at −80°C for 1 hour following removal of the culture supernatant. The culture block was then equilibrated to room temperature and 500 µL 1xPBS added to each well followed by shaking at 1500 rpm to resuspend the cell pellets and allow for release of VHH from the cells. The culture block was centrifuged for 20 min at 2000xg. Nunc Maxisorp plates were coated with mTfR at 1 µg/mL as described above and blocked with 1% bovine serum albumin (BSA). The clarified VHH supernatants were incubated on the mTfR plates for 1 hour at room temperature. The assay plate was washed and peroxidase conjugated goat anti-alpaca VHH domain specific antibody (#128-035-232, Jackson ImmunoResearch) at 0.8 µg/mL was transferred to the plate and incubated for 1 hour at room temperature. Following a final wash, the assay was developed by addition of tetramethylbenzidine (#T5569, Sigma-Aldrich) and absorbance was measured at 650 nm on a Biotek Synergy 2 plate reader. Clones with absorbance values greater than two standard deviations above background were considered of interest and subsequently sequenced.

### Amyloid beta specific VHH production

Paraschiv et al. previously reported the isolation of amyloid beta binding VHH clones [51]. We selected the sequence for the named Nb3 clone for use in this study. The Nb3 amino acid sequence was imported into SnapGene software (GSL Biotech LLC) and reverse translation performed using preferred codon usage for expression in *E. coli*. Additional sequence, including a (Gly-Gly-Gly-Ser)_3_ between VHH domains, was incorporated for cloning into pHEN2 as a tandem dimer as part of the heterotrimer clones synthesized with TfR binders. To reduce the potential for recombination events, the DNA sequence was manually curated to adjust the codon usage and reduce the frequency of repetitive sequence within the Nb3-Nb3 dimer. The affinity of the Nb3-Nb3 dimer was measured using bio-layer intergerometry as described below.

### Bio-layer interferometry (BLI) assessment of VHH binding kinetics

The binding kinetics of the selected VHH clone against mTfR and Aβ was assessed by BLI. For measurements of mTfR kinetics, biotinylated VHH was diluted into assay buffer at 1 µg/mL and immobilized onto streptavidin coated biosensors (#18-5019, Sartorius) to a minimum response value of 1 nm on the Octet Red96 System (Sartorius). For measurements of amyloid beta, beta-amyloid(1-40)-Lys(biotin-LC) (AS-23517, Anaspec), was diluted into assay buffer at 1 µg/mL and immobilized onto streptavidin coated biosensors (#18-5019, Sartorius) to a minimum response value of 1 nm on the Octet Red96 System (Sartorius). Purified mTfR or VHH clones were diluted into assay buffer at the specified concentrations. The immobilized antigen biosensors were allowed to associate at 37C° followed by dissociation in the baseline buffer well location. All assays included a background correction condition to allow for sensor normalization. The ForteBio Data Analysis suite was used to normalize the association curves following background subtraction and Savitzky-Golay filtering. Curve fitting was applied using global fitting of the sensor data and a steady state analysis calculated to determine the association and dissociation constants. All assay steps were prepared in Greiner 96-well plates (#655209) in a volume of 300 µL. Assay buffer was defined as 0.1% BSA (w/v) in 1xPBS.

### Synthesis and expression of VHH constructs

Based on the methods described in **Immunopanning and clone screening** and **BLI assessment of VHH binding kinetics** sections, we synthesized several neurotensin-fused VHH monomers with different binding affinities to mouse transferrin receptor. In addition, we produced single polypeptide VHH heterotrimers that consisted of M1_P96H_, a (Gly-Gly-Gly-Ser)_3_ linker, Nb3, a (Gly-Gly-Gly-Ser)_3_ linker, and Nb3 using the method described in **Amyloid beta specific VHH production**. The VHH dimer Nb3-Nb3 binds to amyloid plaques in brain parenchymal of APP/PS1 mice. These constructs were termed “M1_WT_-NT”, “M1_P96H_-NT”, “M1_AA_-NT”, “M1_R100dH_-NT” and “M1_P96H_-triplet”. The VHH naming convention for P96H and R100dH are based on the Kabat nomenclature and refer to the specific amino acid residue modification positions within the VHH sequence [52]. Another neurotensin fusion with a VHH generated against human TfR, which does not bind mouse TfR called “H1-NT,” was used as control. All VHH constructs were designed using SnapGene Software and synthesized (Twist Bioscience) with corresponding restriction endonuclease sites for direct cloning into pHEN2. Sequence confirmed pHEN2 clones with the various constructs were transferred into BL21(DE3) competent *E. coli* cells (C2527I, New England Biolabs). Transformed cells were grown in terrific broth medium containing carbenicillin at 37°C and 300rpm shaking in baffled flasks. Once the culture density reached an optical density equal to 0.6, IPTG was added to a final concentration of 1mM to induce protein expression. For monomer VHH expression the post-induction incubation temperature was maintained at 37C but reduced to 30C for the M1_P96H_-triplet. Following overnight expression, cells were pelleted by centrifugation and VHHs were extracted through osmotic shock and recovery of the periplasmic fraction [49, 53]. Clarified periplasmic fraction was purified using HisPur™ Ni-NTA Resin (88222, Thermo Fisher Scientific) column chromatography. The eluted VHH proteins were further purified by size-exclusion chromatography (SEC) over a Superdex75 10/300 column on an AKTA Pure (Cytiva). Protein purity was assessed by SDS-PAGE on a 10% Bis-Tris MES acrylamide gel and found to be >95% pure.

M1_P96H_-triplets and H1-triplets were fluorescently labeled with using Alexa Fluor™ 647 succinimidyl ester dye to allow fluorescence confocal microscopy of tissue sections. The triplets were incubated with the succinimidyl ester dye for 1 hour in 50mM sodium carbonate buffer, pH 9.6 and purified by desalting using a 5mL HiTrap desalting column in-line with the AKTA Pure. The binding fidelity of the VHH heterotriplet was assessed by ELISA against mTfR. The fluorescence dye labeled M1_P96H_-triplet and H1-triplet were named “M1_P96H_-triplet-647” and “H1-triplet-647”

### Endotoxin removal

Removal of contaminating endotoxin was achieved using High-Capacity Endotoxin Removal Resin (88270, Pierce) from the VHH preparations. A volume of 0.25ml endotoxin removal resin was added to 1ml of VHH sample. The VHH and resin were mixed for 2hr at room temperature to allow for absorption of endotoxin. The mixture was centrifuged to pellet the resin and the supernatant collected. A volume of 0.25ml fresh endotoxin removal resin was added to the first-pass solution and mixed for another 2hr at room temperature. Following a final centrifugation to pellet the resin, the solution was collected for endotoxin level testing. Endotoxin levels were measured using the Chromagenic Endotoxin Quant kit (A39552, Pierce) according to the manufacturers protocol to ensure a level of endotoxin <0.5 endotoxin units/mg of total protein.

### Binding assessment of histidine mutations by ELISA

To determine the effect on histidine mutation introduction, an ELISA was performed with the post VHH binding wash buffer at normal and reduced pH. Nunc Maxisorp plates were coated with mTfR at 1 µg/mL in 50mM sodium carbonate, pH 9.6 overnight at 4°C. The plates were blocked with 1% BSA in 1xPBS buffer and then triplicate dilutions of each VHH prepared in 0.5% BSA in 1xPBS buffer and transferred to the plate. Binding at room temperature for 2 hours was followed by three cycles of 5-minute washes with either 1xPBS, pH 7.2 or 1xPBS, pH 5.5. Following the pH dependent wash step, the bound VHH was detected with peroxidase conjugated goat anti-alpaca VHH domain specific antibody (#128-035-232, Jackson ImmunoResearch) at 0.8 µg/mL incubated for 1 hour at room temperature.

Following a final wash, the assay was developed by addition of tetramethylbenzidine (#T5569, Sigma-Aldrich), the reaction was terminated by addition of 50 µL 1M hydrochloric acid and absorbance was measured at 450 nm on a Biotek Synergy 2 plate reader.

### Animals

All animal experiments were conducted under protocol approved by the National Institute of Neurological Disorders and Stroke (NINDS)/ National Institute on Deafness and Other Communication Disorders (NIDCD) Animal Care and Use Committee in the National Institutes of Health (NIH) Clinical Center (Protocol Number: 1406-21). C57BL/6J female mice were purchased from Jackson labs at 6-7 weeks of age and used at 7-12 weeks of age. Fifteen mice were divided into five groups with three mice in each group to test the five VHH-NT fusions. Anesthetized mice were injected via tail vein with VHH-NT fusions: M1_WT_-NT, M1_P96H_-NT, M1_AA_-NT, M1_R100dH_-NT and H1-NT for screening. Experiments were performed at the same time each day. To confirm the result from the five VHH-NT fusions, a blinded replication experiment was performed with additional fifteen mice randomized using a random number generator into five groups with three mice in each group. Six mice were randomly assigned into two groups to test the ability of VHH triplet-NT to get across the BBB after fused to anti-Aβ dimer Nb3-Nb3. Three mice were injected with M1_P96H_-Triplet-NT and three mice were injected with H1-Triplet-NT.

APP_SWE_/PSEN1dE9 (MMRRC Strain #034832) positive transgenic mice were purchased from Jackson labs [54, 55], maintained on a hybrid (C57BL/6xC3H) background, and raised under protocols approved by the National Institute of Neurological Disorders and Stroke (NINDS)/ National Institute on Deafness and Other Communication Disorders (NIDCD) Animal Care and Use Committee in the National Institutes of Health (NIH) Clinical Center. Ten transgenic mice ages between 13-15 months, eight males and two females, were randomly assigned to two groups. Five mice were injected with M1_P96H_-triplet-647 and five mice were injected with H1-triplet-647.

### Injection of VHH-NTs and VHH-triplet-647s

Mice were anesthetized with 60% oxygen/ 40% medical air gas mixture containing 5% isoflurane in an induction box. After a stable anesthesia plane was established, mice were maintained at 1.5-2% isoflurane level. Artificial tears ointment was applied to prevent eye injury due to drying. Mice were placed on an electrical heating pad to maintain body temperature at 37C°. The VHH-NTs, M1_WT_-NT, M1_P96H_-NT, M1_AA_-NT, M1_R100dH_-NT and H1-NT were injected into wild-type C57BL/6J mice through the tail vein using a 30 Gauge needle in a single bolus at a dose of 600nmol/kg body weight in 1xPBS. The VHH-triplet-647s, M1_P96H_-triplet-647 and H1-triplet-647, were injected intravenously through single bolus injection at a dose of 1000nmol/kg body weight. Mice were maintained under anesthesia for approximately 2-3 minutes. Following the procedure, the mice were allowed to recover on a heating pad until fully ambulatory and then returned to their home cage with immediate access to food and water.

### Temperature measurement

Infrared thermometry was used for temperature measurement using an infrared thermometer (Model# 62 MAX+, Fluke). Abdominal fur was removed by application of topical depilatory cream prior to temperature measurement for precise data collection. Baseline temperature was measured three times before intravenous (IV) injection with a time interval of 30min between measurements. Then, mouse temperatures were measured at the time intervals of 30min, 1hr, 1.5hr, 2hr, 2.5hr, 3hr, 3.5hr, 4hr, 4.5hr and 5hr after injection. All temperature measurements were performed by investigators blinded to the identity of the injected VHH sample. Mice were briefly anesthetized (<20 seconds) with isoflurane for each temperature measurement. In preliminary experiments we confirmed that this brief anesthesia did not affect temperature.

### Confocal microscopy of mice brains injected with triplet-647

Two hours post IV injection mice were sacrificed under isoflurane anesthesia by cardiac perfusion with 1X PBS + Heparin (10 units Heparin per milliliter 1X PBS). Following perfusion, mice were decapitated using a pair of sharp surgical shears and the brain was carefully excised from the cranium. Brains were fixed in 4% PFA for 24hrs then equilibrated in 30% sucrose for 48 hours. Then, mouse brains were sectioned at 50 µm thickness using a freezing sliding microtome. Staining was performed to visualize amyloid plaques using the Congo Red derivative X34.[56] Tissue was rinsed with 1xPBS two times. Then, tissues were incubated in 40%EtOH/60%PBS at pH10 containing 10 µM X34 (SML1954-5MG, Sigma) for 10min. After X34 incubation, tissue was rinsed with milliQ water five times then differentiated in 40%EtOH/60%PBS at pH10 for 2min. After differentiation, the tissue was rinsed with miliQ water for 10min then mounted onto positively charged slides (EF15978Z, Daigger®). The mounted tissue sections were allowed to dry overnight at room temperature and cover slipped using fluoromount-G (00-4958-02, Invitrogen™).

To objectively evaluate the effect of intravenous injection of M1_P96H_-Triplet-647 and H1-Triplet-647 on APP/PS1 mice, brain sections at around 2.5mm posterior to bregma were selected for confocal fluorescence imaging. The brain sections were equally divided into 10 parts in both horizontal (x) and vertical (y) directions (**S1 Fig**). X and y coordinates were randomly selected using by random numbers from 1 to 10. When (x, y) coordinates fell onto a cortical area, stacks of images were taken which covered 16µm depth (z direction) starting from the top of the brain section. When (x, y) coordinates fell outside of the cortex, the microscope was moved to the next randomized (x, y) coordinates without taking images. Brain sections were imaged using a Zeiss LSM 510 microscope. Stacks of images were acquired with eight images per stack and 2µm optical thickness at 20x magnification using a Zeiss Plan-Apochromat 20x/0.8NA lens. Images were acquired using laser wavelength at 633nm to visualize the VHH triplets conjugated with 647 dye and 405 nm wavelength laser to visualize the X-34 dye. For 647 channel images, laser at wavelength 633nm was used with a LP650 filter and laser power set to be at transmission 100%. For X34 channel images, laser at wavelength 405nm was used with a BP420-480 filter and laser power set to be at transmission 15%. The images were exported in lsm file format for image analysis using ImageJ.

### Fluorescent confocal microscopy - Naïve brain as controls

Naïve APP/PS1 positive mouse brain sections stained only with X-34 for fluorescence confocal microscopy were used as a negative control. Naïve APP/PS1 positive mouse brain sections stained with both X34 and *ex vivo* M1_P96H_-Triplet-647 for fluorescence confocal microscopy were used as a positive control. For *ex vivo* M1_P96H_-Triplet-647 staining, naïve brains were sectioned and washed in 1XPBS. Sections were incubated in 3% normal donkey serum for 30min to block non-specific binding. After blocking, tissue sections were incubated overnight at 4C° in M1_P96H_-Triplet-647 at a concentration of 10ug/ml in blocking solution. Tissues were then washed in 1xPBS three times followed by X34 staining as described.

### Automated image analysis

To quantitatively analyze the confocal microscope images, thresholding and particle analysis were performed to remove background signals and isolate target structures of interest. The percent area of X34 stained amyloid plaques was quantified with thresholding at (0, 30) followed by Particle analysis. To remove low signal background, 647 channel image thresholding was set at 0-1000. Then the 647 channel images was analyzed with particle analysis. The parameters for particle analysis were set to be 0-1000 pixels for size and 0.1-1.0 for circularity. The parameters for thresholding and particle analysis were determined through testing of different parameters to capture the qualitative morphology of amyloid plaques as assessed by experienced investigators. Amyloid plaque levels were represented as % area of X34 coverage and VHH triplet-647 entry into the brain parenchyma was represented as % area of Alexa 647 dye coverage in the areas with X34 staining. Because there is mouse-to-mouse variability in plaque size and X34 staining intensity, we analyzed the ratio of the Triplet-647 coverage to X34 coverage.

### Statistical analysis

Unpaired student t-test were performed to evaluate the differences between the transcytosis ability of M1_P96H_-triplet-647 and H1-triplet-647 to amyloid plaques in APP/PS1 transgenic mice brains. Graphs were created using Prism. Sample sizes were based on availability of transgenic mice and previous experiments. No formal power calculations were performed.

## Results

### Endotoxin removal

VHHs contamination by endotoxins is a byproduct of expression in *E. Coli* cells [57]. Endotoxin can induce systemic inflammation and cause disruptive BBB changes [58]. It was found that the excess endotoxin level in VHH-NT fusions can cause additional hypothermia (**S2 Fig**). To avoid potential hypothermia effects caused by endotoxin, endotoxin was removed after purification. After endotoxin removal, the level of endotoxin level in VHHs was less than 0.1 endotoxin unit/ml (EU/ml) within normal range [59].

### Anti-mTfR VHH variant screening based on binding affinity measurement

VHH monomer-NT (M1-NT and H1-NT) and VHH-heterotrimer-NT (M1_P96H_-triplet-NT) had characteristics consistent with expectation. **S3 Fig and S4 Fig** showed the SDS-PAGE gel and exemplar size-exclusion column results of the VHH monomer-NT and VHH-heterotrimer-NT after purification. VHH monomer-NTs had size around 14kDa (**S3 Fig**) and VHH heterotrimer-NTs had size between 38-49kDa (**S4 Fig**).

Histidine mutations could potentially impart pH dependence of binding [60]. Histidine protonation at lower pH-values can increase dissociation rate of antibodies to their receptors. Maeda et al. found that the dissociation rate was less rapid in the intracellular acidic compartments once histidine was deleted from human epidermal growth factor [61]. In this study, we generated different M1 variants based on the effect of histidine on VHH dissociation rate. **S5 Fig** shows the mTfR ELISA on M1 variants: M1_P96H_, M1_WT_ and M1_R100dH_. Comparing M1 variant binding to mTfR under normal physiological pH (pH = 7.2) and acidic pH (pH = 5.5), lower pH did not alter the dissociation of M1_WT_ to mTfR but substantially impacted the dissociation of M1_P96H_ to mTfR.

Then the affinity of M1 variants to mTfR was measured using Octet. Based on Octet measurement, the affinity (KD) of M1_WT_-NT, M1_P96H_-NT, and M1_R100dH_-NT to mTfR were <1nM, 1.12nM, 1.52nM. While the affinity (KD) of M1_AA_-NT and H1-NT to mTfR was not detectable. **Fig 2** shows the association and

**Fig 2.**
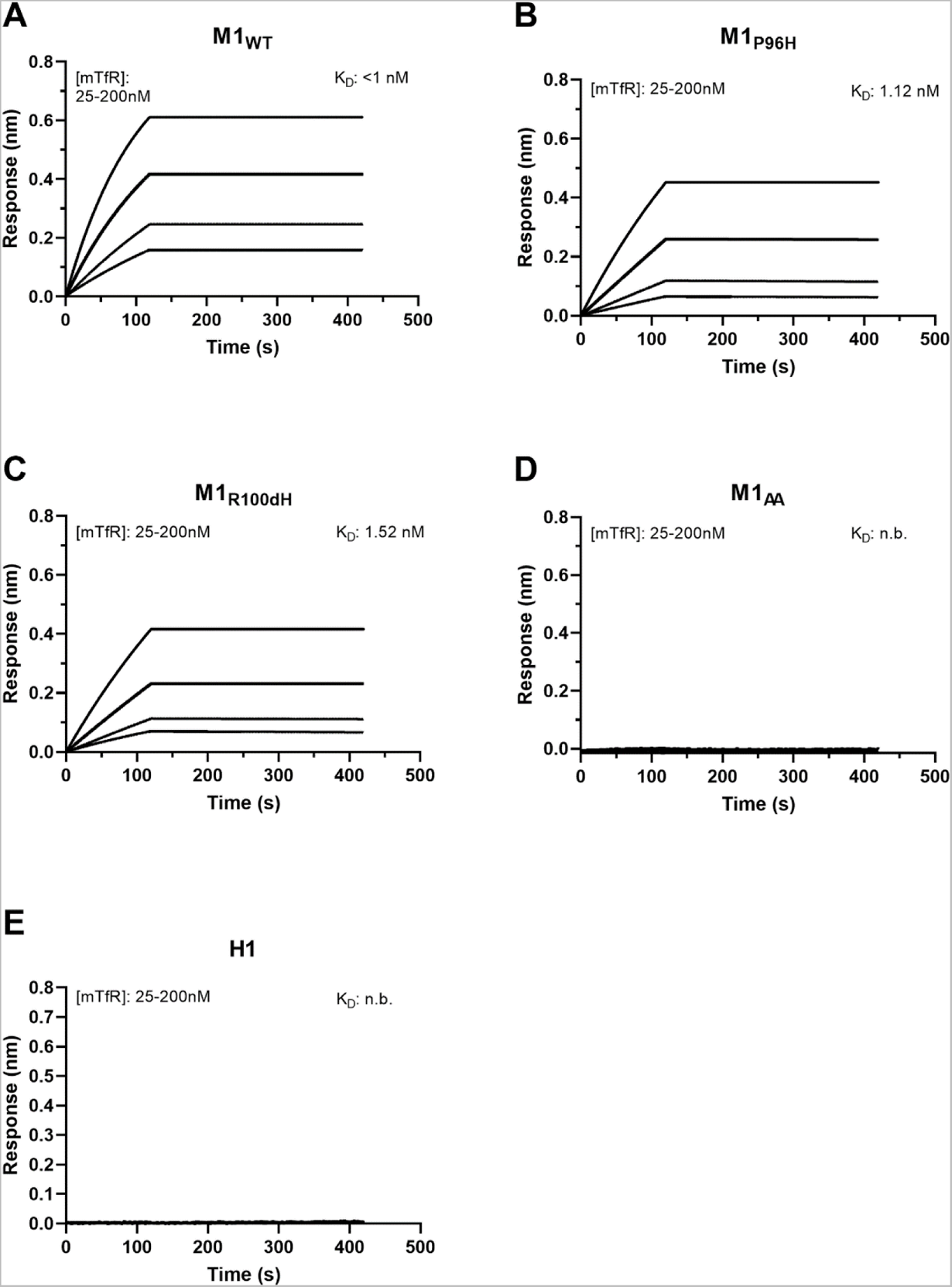
Affinity binding curves of the five M1-NT variants. M1_WT_-NT (**A**), M1_P96H_-NT (**B**), M1_R100dH_-NT (**C**), M1_AA_-NT (**D**) and H1-NT (**E**) binding affinity (K_D_) to mTfR. Using biolayer interferometry on an Octet Red96 system, association and dissociation rates were determined by immobilizing biotinylated VHH onto streptavidin-coated optical sensors. The M1-NT variant association and dissociation curves to mTfR were plotted and the affinity of each to mTfR was calculated. (n.b. = no binding) dissociation curves of the M1_WT_-NT, M1_P96H_-NT, M1_AA_-NT, M1_R100dH_-NT and H1-NT to mTfR. M1_WT_-NT, M1_P96H_-NT, and M1_R100dH_-NT bond to mTfR well while M1_AA_-NT and H1-NT showed no binding. M1_WT_-NT and M1_P96H_-NT showed no binding to human TfR (data not shown).

### M1 variant screening for BBB transcytosis using neurotensin fusion and hypothermia assessment

To assess BBB transcytosis ability, several M1 variants were fused to NT and screened based on the extent of hypothermia effects. Three M1 variant NT fusions, including M1_WT_-NT, M1_R100dH_-NT and M1_P96H_-NT were injected to WT mice. NT alone was injected into WT mice at the same molarity as an initial negative control. At a dose of 600nmol/kg body weight, the M1 variants with different binding affinities to mTfR show different hypothermia effects (**Fig 3**). M1_WT_-NT reduced temperatures by approximately 2C°, with an effect that lasted approximately 2 hours. M1_P96H_-NT, at the same dose, decreased temperature by 6C° with effects that lasted more than 4 hours, indicating that M1_P96H_-NT appeared to improve CNS penetration. NT alone did not show any hypothermia effect, indicating that peripherally administered NT alone does not appear to cross the BBB into the CNS. M1_R100dH_-NT gave less than 2C° of hypothermia effect. Mouse body temperatures were stable at baseline.

**Fig 3.**
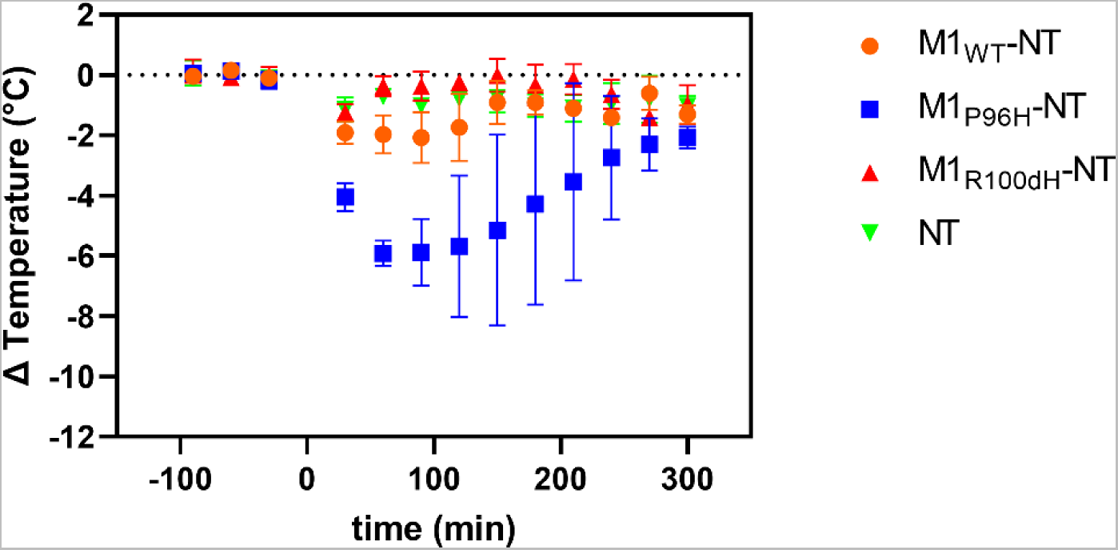
The use of NT fusion to VHHs for VHH screening via hypothermia as an indication of CNS target engagement. The M1 variants with different affinities to mTfR have different hypothermic effects. Among the three M1 variants injected to mice at the same dose 600nmol/kg body weight (n = 3 per group), M1_P96H_-NT produced the most prominent hypothermia effect, with a maximum temperature drop of about 6C° and duration for about 4hrs.

To confirm and extend these findings, a blinded experiment was performed. The previous three M1 variants, plus a fourth mutant M1_AA_-NT that has minimal TfR binding and a different VHH that binds to human TfR called H1 were tested by an investigator blinded to the identity of the injected materials. M1_AA_-NT and H1-NT were tested at a higher molarity in a previous experiment and showed no obvious hypothermia effects (**S6 Fig**). The blinded experiment results were consistent with the previous finding: M1_P96H_-NT clearly reduced the temperature the most, with a maximum drop of about 4C°. The other M1-NTs and H1-NT did not reduce the temperature substantially (**Fig 4**). These results indicate that temperature measurement experiments facilitated the identification of an anti-mTfR VHH variant which induced substantial CNS effects.

**Fig 4.**
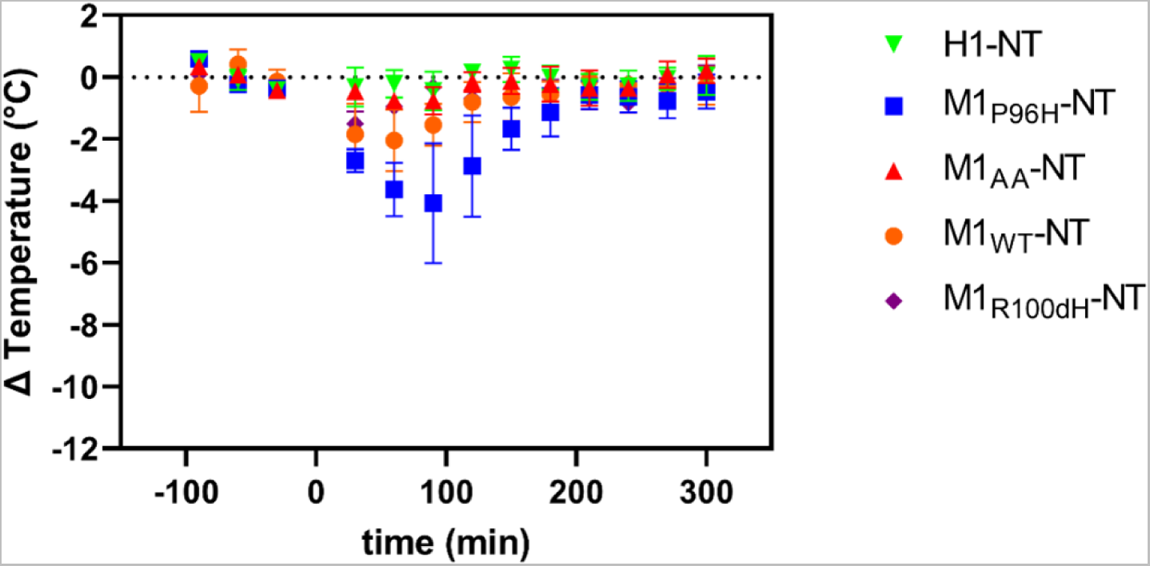
Hypothermic effect of M1 VHH-NT fusions in a randomized, blinded cohort. Five VHH-NT fusions M1_WT_-NT, M1_P96H_-NT, M1_AA_-NT, M1_R100dH_-NT and H1-NT were blinded by an independent investigator. Each blinded VHH-NT fusion was injected in a randomized fashion at a dose of 600nmol/kg body weight (n=3 per group) and body temperature measured over the indicated time interval. Following complete data collection of the temperature measurements, the results were unblinded. Consistent with previous finding, M1_P96H_-NT gave the most prominent hypothermia effect.

### M1P96H-NT dose effect

To understand whether the amount of M1_P96H_-NT injected to mice affects the extent of hypothermia effects, we injected M1_P96H_-NT to WT mice at four different doses: 67nmol/kg body weight, 200nmol/kg body weight, 600nmol/kg body weight, and 1800nmol/kg body weight. Twelve mice were used in this experiment, with three mice at each dose (**Fig 5**). At the two lower doses, there was minimal hypothermia effects. With increased M1_P96H_-NT concentration, the maximum temperature drop increased to about 8C° and also the duration of hypothermia was extended to as long as 5 hours. Thus, these was an apparently monotonic dose-response relationship between intravenous M1_P96H_-NT and hypothermia.

**Fig 5.**
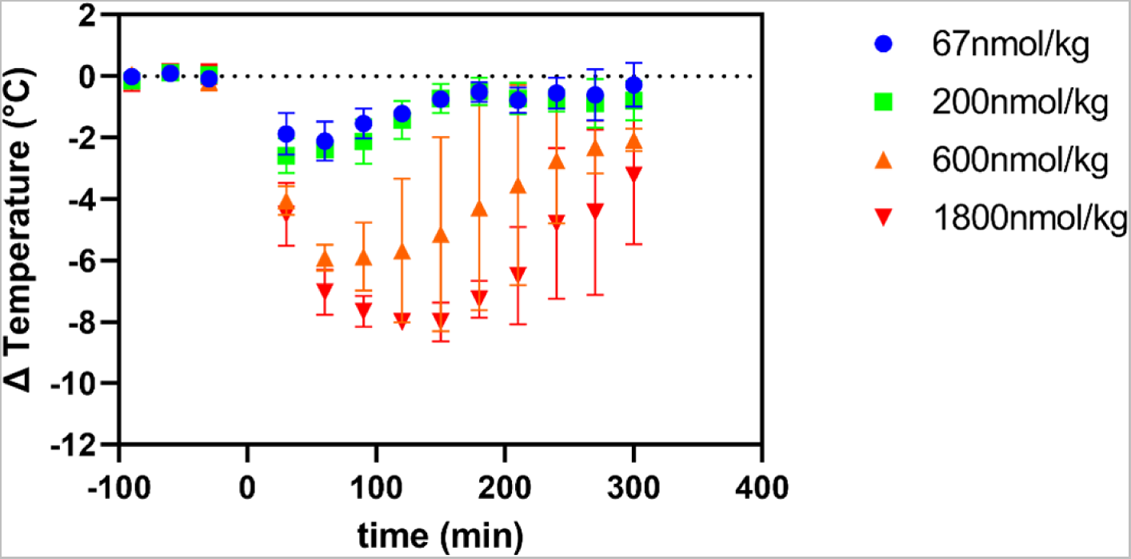
M1_P96H_-NT dose effect in WT mice after IV injection. This figure shows the dose effect of M1_P96H_-NT in WT mice after tail vein injection (n = 3 per group). The hypothermia effect was stronger and lasted longer time with dose increased from 67nmol/kg body weight to 1800nmol/kg body weight.

### Temperature effect of M1P96H-Triplet-NT and H1-Triplet-NT

Based on the octet results and temperature measurement, M1_P96H_ was chosen as the variant to use to assess ability to cross the BBB after conjugating to a payload. As a proof of concept, we used the anti-Aβ VHH dimer Nb3-Nb3 as the payload fused in a single polypeptide with M1_P96H_ and NT and named M1_P96H_-Triplet-NT. The Nb3-Nb3 dimer was also fused to H1 and NT and named H1-Triplet-NT as a negative control. The M1_P96H_-Triplet-NT and H1-Triplet-NT were injected intravenously into six WT mice (three mice for each Triplet-NT fusion). Body temperature was measured as described in the Temperature measurement section. **Fig 6** shows the temperature change after tail vein injection of M1_P96H_-Triplet-NT and H1-Triplet-NT. For M1_P96H_-Triplet-NT, there was about 4C° temperature drop after the intravenous injection and the hypothermia effect peaked at about 1 hour post injection. There was no substantial hypothermia effect after intravenous injection of H1-Triplet-NT. This result confirmed that after fusing to an anti-Aβ VHH dimer payload, the M1_P96H_-Triplet-NT was still able to get across the BBB and exert a CNS effect.

**Fig 6.**
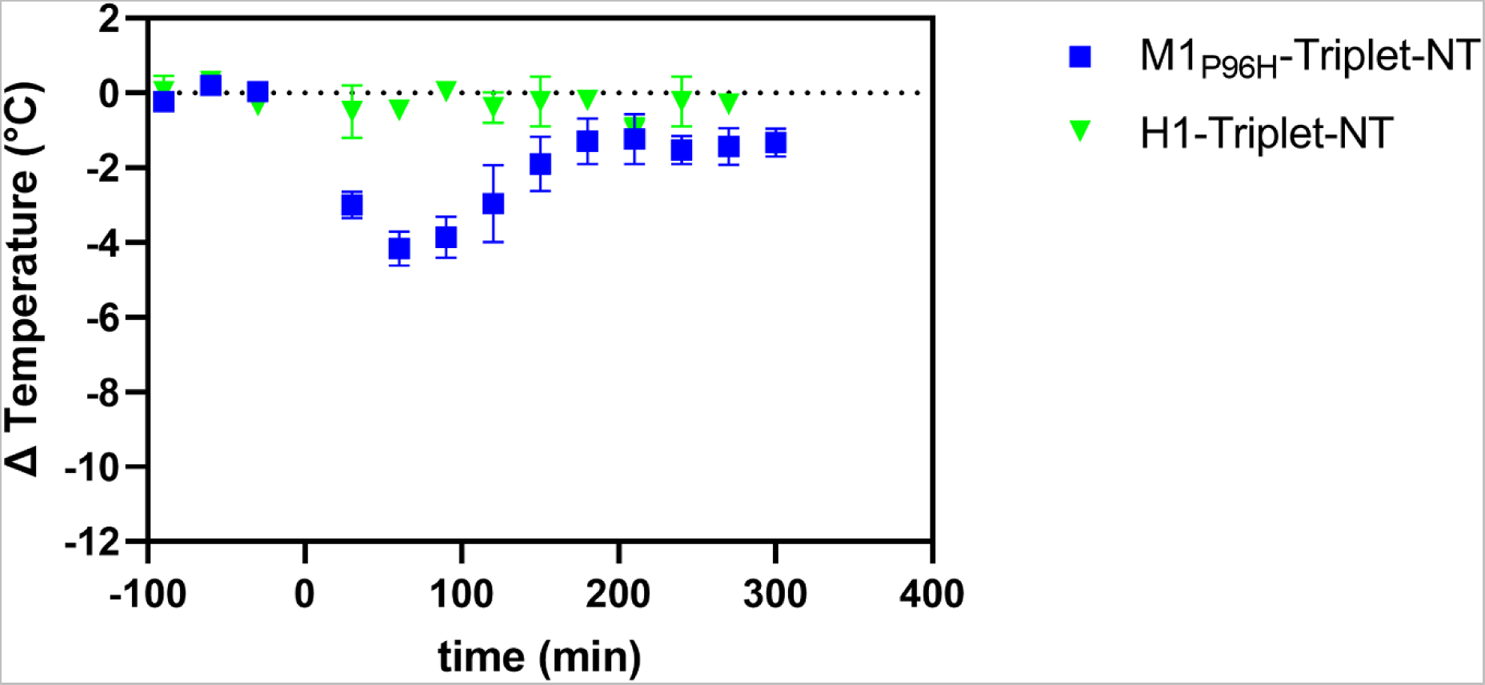
Measurement of M1_P96H_-Triplet-NT and H1-Triplet-NT hypothermia effect. M1_P96H_-Triplet-NT and H1-Triplet-NT were injected at dose of 600nmol/kg body weight (n = 3). M1_P96H_-Triplet-NT caused a temperature drop of about 4C° and H1-Triplet-NT caused no temperature drop.

### Affinity measurement of VHH-Triplet

To confirm that fusion with Nb3-Nb3 did not affect the binding ability of M1_P96H_ and Nb3-Nb3 dimer to their targets, the affinity of M1_P96H_-triplet to mTfR and to Aβ were measured to be 1.63nM and 9.98nM using Octet (**S7 Fig**). The affinity of the M1_P96H_-triplet to mTfR was on the same order before and after the fusion. The affinity of H1-Triplet-647 to mTfR and Aβ was also assessed using Octet and measured to be no binding to mTfR and 9.44nM to Aβ (**S7 Fig**). It was verified that the fusion of M1_P96H_ and H1 to VHH dimer Nb3-Nb3 did not change their binding affinities.

To ensure that the dye labeling did not affect M1_P96H_-triplet binding to mTfR, a mTfR ELISA on M1_P96H_-triplet before and after Alexa 647 dye labelling was performed. The binding curves before and after labeling overlapped with each other (**S8 Fig**), indicating that the Alexa 647 dye labeling did not affect the binding.

### Direct assessment of brain target engagement after intravenous injection of VHHs using confocal microscopy

VHH-Triplets were injected to APP/PS1 positive transgenic mice to directly assess brain target (Aβ) engagement. Ten transgenic mice were randomized into two groups, with five mice in each group. Five mice were injected with M1_P96H_-Triplet-647 and five mice were injected with H1-Triplet-647 as a negative control which does not bind to mTfR. Because the hypothermia effect was most prominent in the first two hours post injection in previous experiments using NT fusions, transgenic mice were sacrificed two hours post IV injection. **Fig 7** shows exemplar confocal images of brains injected with M1_P96H_-triplet-647 or H1-triplet-647. From the exemplar images, it appeared that most of the 647 signals were similar to background, with some areas that had morphology consistent with amyloid plaque labeling. However, surprisingly, the confocal imaging patterns were similar in both M1_P96H_-triplet-647 and H1-triplet-647 injected mice, suggesting a modest amount of non-specific BBB crossing and Aβ plaque binding plus background autofluorescence, rather than the expected extensive mTfR-mediated transcytosis and Aβ plaque binding.

**Fig 7.**
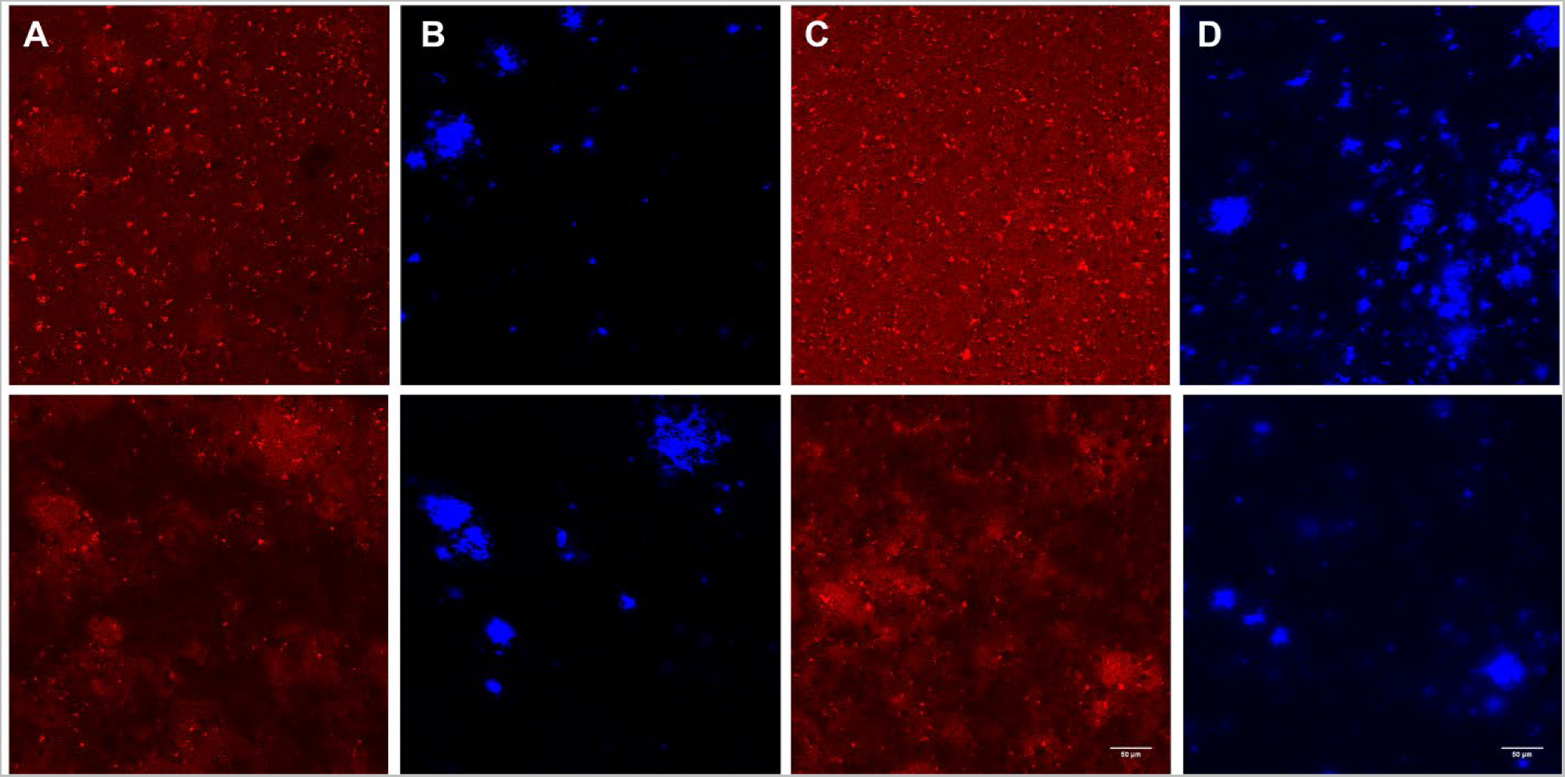
Representative immunofluorescent confocal microscopy of mice brains 2 hours after intravenous injection. 647 channel (**A**) and X34 channel (**B**) microscopy of mice brains injected with H1-Triplet-647. 647 channel (**C**) and X34 channel (**D**) microscopy of mice injected with M1_P96H_-Triplet-647.

To confirm that the lack of anti-Aβ plaque binding after IV injection by M1_P96H_-triplet-647 was not caused by the inability of the Nb3-Nb3 to label Aβ in the context of the M1_P96H_-triplet-647 fusion construct, an *ex vivo* labeling experiment was performed. We labeled brain sections from a naïve APP/PS1 positive mouse with M1_P96H_-Triplet-647 using modest concentrations and no antigen retrieval. **Fig 8** shows that the Nb3-Nb3 dimer part of M1_P96H_-Triplet-647 was able to bind to amyloid plaques, with substantially enhanced fluorescence signals in areas of plaques. Thus, the modest plaque labeling after intravenous injection with M1_P96H_-triplet-647 is not likely to be attributed to lack of plaque binding affinity. Autofluorescence was relatively modest in the 647 channel in sections labeled with only X34 (**Fig 8C, D**).

**Fig 8.**
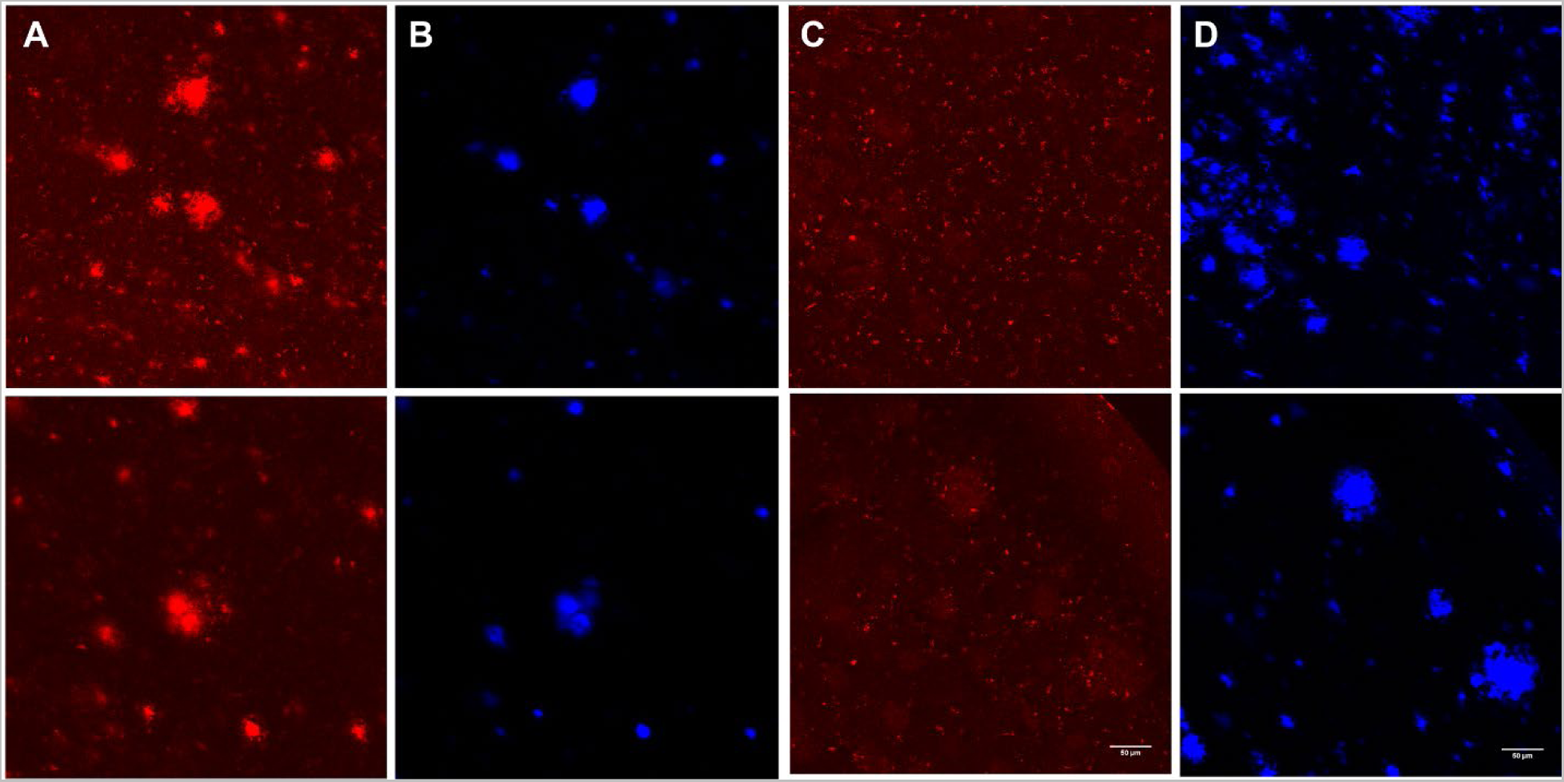
Representative immunofluorescent confocal microscopy of ex vivo APP/PS1 positive mouse brain sections. 647 channel (**A**) and X34 channel (**B**) microscopy of the ex vivo brain sections labeled with M1_P96H_-Triplet-647. 647 channel (**C**) and X34 channel (**D**) microscopy of the ex vivo brain sections without M1_P96H_-Triplet-647 labeling.

Because of the possibility of subconscious bias in selecting regions for imaging and to assess for more subtle quantitative differences between M1_P96H_-triplet-647 and H1-triplet-647 plaque labeling, we performed randomized, blinded, automated analyses of the brain sections to objectively compare the confocal imaging results. Eleven or twelve confocal stacks of images were acquired at randomly selected x, y coordinates in cortex (**S1 Fig**). Confocal images of brain sections were processed using ImageJ for quantitative assessment.

The % area of X34 coverage, % area of Triplet-647 coverage and the % area ratio of Triplet-647/X34 coverage were averaged across the eleven to twelve images stacks acquired for each mouse brain. The average values of % area of each mouse brain was plotted in **Fig 9** Comparing mouse brains injected with M1_P96H_-triplet-647 or H1-triplet-647, the average values of % area of Triplet-647 coverage/% area X34 coverage had no significant difference (**Fig 9C**). There was also no significant difference for % area X34 coverage or % area Triplet-647 coverage (**Fig 9A, B**).

**Fig 9.**
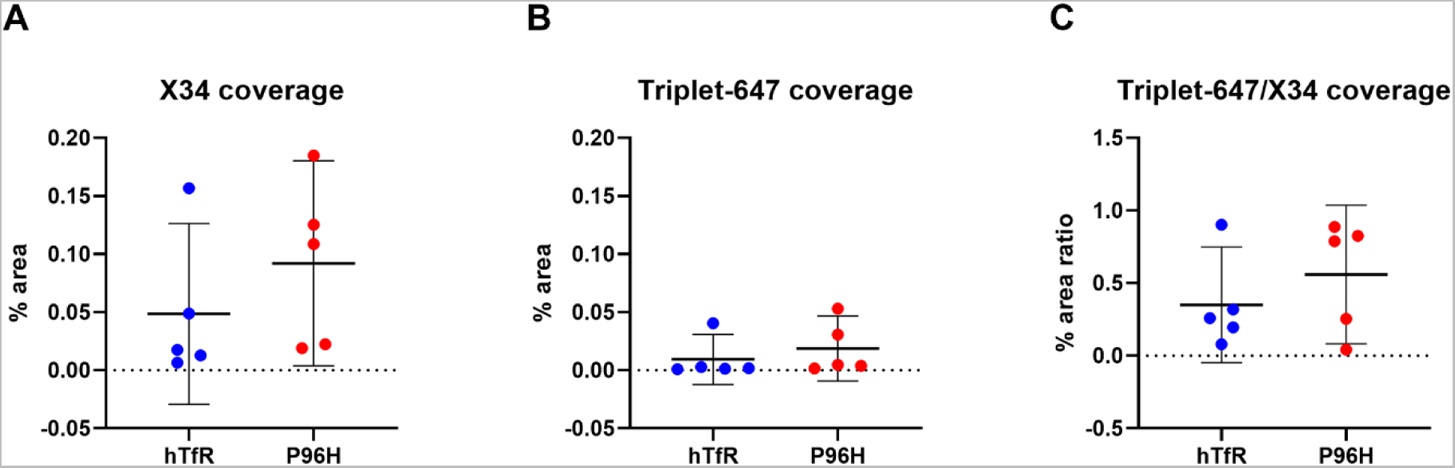
% area and % area ratio of X34 channel images and 647 channel images. (**A**) The % area of X34 coverage in the X34 images. There was no significant difference for mice injected with H1-triplet-647 or M1_P96H_-triplet-647; unpaired student t-test, p= 0.3341 (**B**) The % area of Triplet-647 coverage in 647 channel images. There was no significant difference for mice injected with H1-triplet-647 or M1_P96H_-triplet-647; unpaired student t-test, p = 0.4815. (**C**) The % area ratio of Triplet-647/X34 coverage. There was no significant difference for mice injected with H1-triplet-647 or M1_P96H_-triplet-647; unpaired student t-test, p = 0.3788. 647 channel images and the ratio of 647 channel/X34 channel images.

### Alternative Confocal Image Processing – thresholding and intensity measurement

To ensure the finding was not affected by the specific analysis method, confocal images of brain sections were analyzed in several alternative ways using ImageJ (**S9 Fig**). First, X34 images were alternatively thresholded at (0, 10) and (0, 30) and were converted to binary images. Because amyloid-beta pathology can extend beyond the boundaries of X34 positive fibrillar plaque cores, the X34 images thresholded at (0, 30) were also processed with dilation in ImageJ to add pixels from the edges of plaques. These plaque and peri-plaque regions identified on the X34 images were applied to the 647 channel images to calculate the mean 647 signal of these regions. The mean 647 signals were averaged across image stacks acquired for each mouse brain The average values of mean 647 signals of each brain were plotted in **Fig 10** There was no significant difference of the mean 647 signal between the two groups in any of these analyses. These findings confirmed that there was no detectible receptor mediated transcytosis of M1_P96H_-triplet-647 into the cortex, despite good evidence for a CNS effect for the same construct when fused to neurotensin.

**Fig 10.**
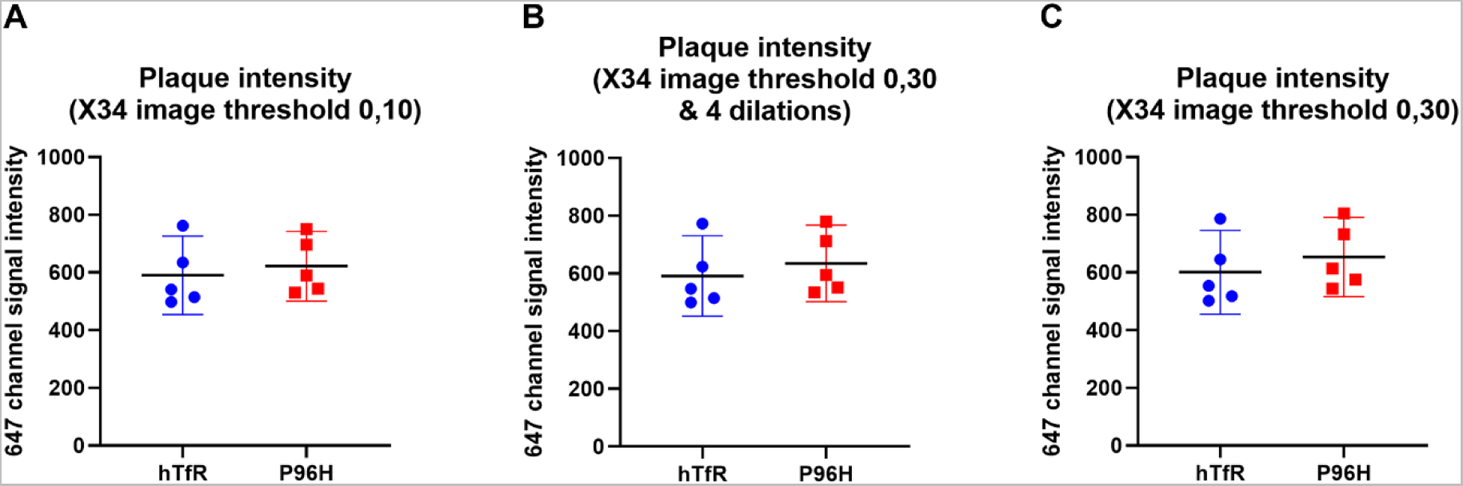
Averaged mean fluorescence in the 647 channel of plaques containing regions. (**A**) The threshold for X34 images was set to be at (0, 10) There was no significant difference between the mean signals from mice injected with H1-triplet-647 and mice injected with M1_P96H_-triplet-647; unpaired student t-test, p = 0.6404. (**B**) The threshold for X34 images was set to be at (0, 30), then the image was eroded for four times. There was no significant difference between the mean signals from mice injected with H1-triplet-647 and mice injected with M1_P96H_-triplet-647; unpaired student t-test, p = 0.5527. (**C**) The threshold for X34 images was set to be at (0, 30). There was no significant difference between the mean signals from mice injected with H1-triplet-647 and mice injected with M1_P96H_-triplet-647; unpaired student t-test, p = 0.4829.

## Discussion

In this study, a modular system was designed to efficiently test the ability of anti-mTfR VHHs to cross the BBB using NT-induced hypothermia as a readout. Because only NT in the CNS can induce hypothermia effects [39, 46, 48, 62], experimentally measured hypothermia was used to infer the VHHs’ ability to cross the BBB through transferrin receptor mediated transcytosis followed by VHH-NT fusion binding to neurotensin receptors. Using this NT-based screening system, this study successfully identified an anti-mTfR VHH variant, M1_P96H_, which has good binding properties to the mouse TfR and appeared to mediate BBB transport efficiently. To attempt to validate the ability of this anti-mTfR VHH to carry cargos across BBB, this anti-mTfR VHH was fused with an anti-Aβ VHH dimer and fused to NT. The H1 VHH was fused with anti-Aβ VHH dimer and NT as negative control. The M1_P96H_-Triplet-NT retained substantial hypothermia effects while the H1-Triplet-NT showed no hypothermia effect after intravenous injection. This finding confirmed the BBB transcytosis ability of M1_P96H_-Triplet-NT. However, the modular system for VHH screening does not appear to translate effectively into target binding in the cortex. To assess brain target engagement, M1_P96H_-triplet was conjugated to the fluorescent Alexa 647 dye and the dye conjugates were injected intravenously into APP/PS1 positive transgenic mice. A similar VHH triplet dye conjugate, H1-triplet which does not binding mTfR was used as a control. Surprisingly, there was no significant difference of the amyloid plaque binding between these two VHH triplet dye conjugates. This result was not likely due to failure of plaque binding, because the VHH triplet dye conjugates bound plaques well in *ex vivo* experiments. Thus, the likely explanation for our findings was that the VHH triplet dye conjugates did not effectively cross the BBB in the cortex; the hypothermia effect induced by NT did not correspond to target engagement in cortex. The finding indicates that the use of NT as a rapid screening platform for anti-mTfR VHHs does not necessarily predict generalized cargo delivery across the BBB.

The lack of correspondence between NT induced hypothermia after intravenous injection and brain target engagement could be possibly explained by the concept that the BBB is not a unitary phenomenon, and the permeability of BBB may be different from brain region to brain region. The hypothalamus is the center for thermoregulation, and likely to be the site of action of NT in the brain. The preoptic area of hypothalamus contains microcircuitry through which cutaneous and core thermal signals are integrated for thermoregulation and temperature homeostasis [63, 64]. Studies have found that the interface between the median eminence and the arcuate nucleus of the hypothalamus is somewhat leaky to molecules in the circulation [65, 66]. Cheunsuang et al. confirmed the specialized nature of median eminence and medial arcuate nucleus BBB by testing the distribution of hydroxystilbamidine and wheat germ agglutinin after intravenous injection [65]. They found that hydroxystilbamidine was taken into the median eminence and medial part of the arcuate nucleus, while the wheat germ agglutinin diffusely distributed in the arcuate nucleus and median eminence following intravenous injection. Furthermore, Morita and Miyata reported the accessibility of low molecular weight blood-derived molecules to the parenchyma in the median eminence and arcuate nucleus [67]. The hypothermia effect caused by NT binding to NT receptor is likely to be mediated in the hypothalamus [46]. Young et al. explored the distribution of NT receptors in rat brains and found moderate to high density of NT receptors in hypothalamus. The hypothermia effect was induced when NT was injected to medial, lateral preoptic and anterior area of the hypothalamus. Thus, the dissociation between robust NT-mediated hypothermia effects vs. negligible binding to amyloid plaques in cortex after IV injection could be explained by a relatively leaky hypothalamic BBB. Future studies will be required to directly test this possibility.

Also, in this study, the minimum required dose for NT induced temperature effect was about 20-fold higher than Stocki et al. [39]. The relationship between the amount of VHHs needed for NT-induced hypothermia CNS effect and the amount of VHHs required to label and visualize the amyloid plaques is unclear. Meanwhile, the affinity of the anti-Aβ VHH Nb3-Nb3 was measured to be around 9-10nM, which is about 10-fold lower than the affinity of anti-TfR VHH M1_P96H_. Higher affinity Aβ binding antibodies would be required to assess the role of differential affinity for Aβ vs TfR. Because TfR is also expressed on other brain cells such as choroid plexus epithelial cells and neurons [68], it is possible that the M1_P96H_-Triplet-NT crosses the blood CSF barrier and engages with periventricular hypothalamic NT receptors more effectively than it cross the blood brain barrier to interact with parenchymal Aβ. Anti-TfR antibodies such as OX-26 and 8D3 could be tested as a positive control to explore the correlation between NT induced hypothermia and brain target (Aβ) engagement in the future [19, 21]. The absolute amount of anti-TfR positive controls and VHH-Triplet in the brain parenchyma after crossing the BBB will be accessed using ELISA after separating brain capillaries and brain parenchyma [39, 69].

### Limitations

There are several limitations of this study. First, although the M1_P96H_-NT induced substantial hypothermia, the kinetics of this anti-mTfR VHH variant have not been fully optimized. **Fig 2** shows that the dissociation rate of the M1_P96H_ from mTfR was 1.33×10^−4^ which could be still too low. Based on Hultqvist et al., after the VHH binds to the TfR and the complex is internalized, the VHH needs to dissociate from the TfR to get released efficiently into the brain parenchyma [22]. When the dissociation rate is too slow, VHHs may not unbind from the TfR and not get released into the brain parenchyma. Second, it is possible that dye modification with Alexa 647 affected BBB transcytosis even though it did not affect mTfR binding affinity *in vitro* (**S8 Fig**). Alternative labeling strategies will need to be tested in future experiments. Reduction in autofluorescence due to lipofuscin [70] may also improve the sensitivity of fluorescent dye-based detection methods. Third, to match the maximum hypothermia effect, 2hr post injection was chosen as the terminal time point to access brain target engagement. However, this time point may not be optimal. Multiple time points post injection will need to be tested to find the optimal terminal time point for brain target engagement studies. Fourth, the APP/PS1 mice used in the study can also develop astrogliosis [71]. It is possible that the astrogliosis around the plaques could impair BBB transcytosis. Fifth, we do not know the mTfR binding epitope of M1, though it does not appear to interfere with transferrin binding (not shown). Also, it is formally possible, though unlikely, that older APP/PS1 transgenic mice have a less permeable BBB than young WT mice; we have not directly tested hypothermia effects in older APP/PS1 mice. A minor limitation is that the temperature measurement was conducted using an infrared thermometer. Although belly fur was removed to decrease variation and increase the accuracy and precision of temperature measurement, the use of infrared thermometer to measure belly temperature still introduces inter- and intra-user error. Also, the mice need to be anesthetized for acquisition of a steady temperature using the thermometer.

Although we kept the anesthesia time short (about 20 sec) each time to minimize the influence of anesthesia on mouse body temperature. The use of anesthesia still introduced extra variations. Other temperature measurement methods such as the use of implanted thermometers which provide real-time temperature readings and eliminate interference with mice should be considered for future experiments. Finally, we acknowledge that only one brain section posterior to bregma was analyzed for each mouse. More brain sections for each mouse which cover additional brain areas, including hypothalamus, will need to be imaged and analyzed in the future.

### Future directions

There are many important future directions for this line of investigation. We are in the process of screening other VHH variants for anti-mTfR and anti hTfR-mediated BBB transcytosis. Because the NT induced hypothermia effect does not correspond to target engagement of targets in other brain areas like cortex, an alternative, efficient approach to assess BBB transcytosis is needed. To allow high throughput testing and prediction of therapeutic and diagnostic agent delivery to the CNS, *in vitro* BBB models have been used. Cecchelli et al. developed a human *in vitro* BBB model using cord blood-derived hematopoietic stem cells [72]. This model shows good correlation between *in vitro* predicted ratio of unbound drug concentration in brain and *in vivo* ratio reported in humans. Shayan et al. made a murine *in vitro* BBB model using murine brain microvascular endothelial cells which also shows good correlation of compound permeability compared with *in vivo* values [73]. However, these *in vitro* models do not always reflect the *in vivo* BBB function. Garberg et al. evaluated multiple different *in vitro* models in comparison with an *in vivo* mouse brain uptake assay to understand the *in vitro* models’ potential to predict *in vivo* transport of compounds across BBB [41]. Low correlations between *in vitro* and *in vivo* data were obtained with a total of twenty-two compounds. Because of the complexity of the *in vivo* environment, *in vitro* BBB models were not recommended to be used in isolation to assess target engagement in brain [74]. There are many other methods to assess BBB transcytosis. For example, Stocki et al. tested the BBB transcytosis of a variable domain of new antigen receptors (VNAR) TXB2 by fractionating capillaries from the brain parenchyma and measured the concentration in capillaries and brain parenchyma [39]. Yu et al. assessed the uptake of an anti-TfR antibody by homogenizing the brain target areas and measuring the antibody concentration with ELISA [17]. A new screening platform will need to be established and tested in the future to facilitate of the discovery of optimal anti-mTfR VHHs.

In this study, the VHHs were produced in *E. Coli.* and contaminated by endotoxins in the outer membrane of this gram-negative bacteria [57]. Extra steps were required to remove endotoxin from the VHHs. In the future, VHHs will be produced in yeast which has high yield for VHH production and avoid contaminating VHHs with endotoxins [75]. Also, the immunogenicity and toxicity profiles of these foreign protein constructs will be assessed.

Previous research found that anti-TfR antibodies could cause a reduction of reticulocyte count and acute clinical signs [76]. There were no apparent safety concerns raised during the experiments reported here. However, the safety profile of the VHHs will be formally assessed in the future to make sure that the binding of VHHs to the mTfR does not cause reticulocyte reduction or interfere with physiological iron uptake [77].

In addition, extension of VHH fusion protein half-life in the circulation may improve BBB transcytosis. The half-life of single VHHs and VHH triplets was measured to be 2-3 minutes in the mouse after IV injection [78]. The lack of differences in brain signal may be caused by the fast clearance of the VHHs, which did not allow enough VHHs to bind to the TfR on the brain endothelial cells. Conjugation of the VHHs to albumin binding domains, immunoglobulin Fc domains or PEG to slow down VHH clearance and enhance blood residence time will be tested in the future [79–81].

While not directly related to the main aims of the project involving development of brain MRI molecular contrast agents, the mechanisms underlying potential differences in BBB function in the hypothalamus vs. cortex are worthy of further investigation. To begin, target engagement of the Triplet-647 in hypothalamus will need to be assessed. The hypothalamus area of the brain sections will be imaged with confocal microscope and analyzed using ImageJ. Then, the M1_P96H_ could be conjugated to nanobodies against another widely distributed endogenous target such as the ATP-gated ion channel P2X7, which is widely distributed throughout the brain [82]. M1_P96H_-anti P2X7 VHH fusion would then be intravenously injected to mice and its target engagement in cortex and hypothalamus would be assessed and compared. Furthermore, different sizes of dextrans could be intravenously injected to the mice to understand the BBB penetration capacity in hypothalamus and other brain regions [67].

The ultimate goal of this study is to develop a family of molecular MRI contrast agents for diagnosis and assessment of neurodegenerative diseases. Misdiagnosis of neurodegenerative diseases is common because of their heterogeneous nature [83]. Early diagnosis of neurodegenerative diseases could help with early treatments and delay hospitalization, and accurate identification of target populations could help with the development of new treatments [83, 84]. MRI has been widely used for imaging neurodegenerative diseases; however, structural MRI provides indirect and nonspecific measurements. Neurodegenerative diseases are characterized by abnormal accumulation of misfolded proteins including α-synuclein, tau, TDP-43 and Huntingtin in the CNS. Novel families of MRI contrast agents with VHH and IONPs would allow visualization of pathologically specific biomarkers in the living human brain: each VHH would bind to the misfolded proteins while the IONPs provide T1 MRI signals. We have characterized the *in vivo* pharmacokinetics of the contrast agent and optimized the MRI sequence for MR T1 imaging [78, 85], but have not optimized BBB transcytosis of contrast agents. IONPs have been widely in a research context as MRI molecular contrast agents but are not typically used in clinical practice. Liu et al., developed an oligomer-specific scFv antibody W20 conjugated superparamagnetic iron oxide nanoparticles which specifically bound to oligomers in transgenic mouse models of Parkinson’s disease or Huntington’s disease and provided MRI signals [86]. Sillerud et al. synthesized an anti-AβPP conjugated superparamagnetic iron oxide nanoparticle for MRI detection of amyloid plaques in AD [87]. However, none of these contrast agents have provided optimal BBB penetration to allow high quality *in vivo* imaging. The ability of molecular contrast agents to cross the BBB and bind to brain targets in sufficient quantity to give conspicuous MRI signals will need be tested with *in vivo* MR imaging in the future.

## Conclusions

This study used a NT based modular system to screen anti-mTfR VHHs for transferrin receptor mediated transcytosis across BBB. A M1 variant, M1_P96H_, was identified with good performance in inducing hypothermia, an effect which requires crossing the BBB. This M1 variant was fused to the anti-Aβ VHH dimer and labeled with Alexa 647. Surprisingly, however, the dye labelled VHH did not show detectible labeling of amyloid plaques compared with controls after intravenous injection into transgenic mice. Other methods assessing VHH BBB transcytosis will need to be developed for screening VHHs to facilitate the development of MRI molecular contrast agents.

## Acknowledgements

This research was supported by the Intramural Research Program of the NIH, NINDS. We thank Dr. Alan Koretsky for providing a supportive intellectual environment. We thank the NHLBI Biophysics Core Facility for the use of BLI instrument. We thank the NINDS Light Imaging Facility for use of the confocal microscope. We thank Elvira Rodionova and Caroline Francescutti for expressing the VHHs. We thank Dr. Yoshimi Enose-Akahata for assistance with the blinded experiment. The first author would like to thank her PhD thesis committee, Dr. Philip Bayly, Dr. Dennis Barbour, Dr. Hongyu An and Dr. Vijay Sharma for insightful suggestions.

## Disclaimer

The authors have no conflicts of interest to disclose. The views, information or content, and conclusions presented do not necessarily represent the official position or policy, nor should any official endorsement be inferred, on the part of the National Institutes of Health, the Uniformed Services University, the Department of Defense, Henry M. Jackson Foundation for the Advancement of Military Medicine, Inc., or other government agency.

## Availability of data and materials

The datasets used and/or analyzed during the current study are available from the corresponding author on reasonable request.

## Authors’ contributions

**Conceptualization**: Shiran Su, Thomas J. Esparza, David L. Brody.

**Data curation**: Shiran Su, Thomas J. Esparza.

**Formal analysis**: Shiran Su, Thomas J. Esparza.

**Funding acquisition**: David L. Brody.

**Investigation**: Shiran Su, Thomas J. Esparza.

**Methodology**: Shiran Su, Thomas J. Esparza.

**Resources**: Thomas J. Esparza.

**Software**: Shiran Su.

**Writing – original draft**: Shiran Su, David L. Brody.

**Writing – review & editing**: Shiran Su, Thomas J. Esparza, David L. Brody

## Notes

### Competing Interest Statement

The authors have declared no competing interest.

