## Supporting information for "Selection of single domain anti-transferrin receptor antibodies for blood-brain barrier transcytosis using a neurotensin based assay and histological assessment of target engagement in a mouse model of Alzheimer’s related amyloid-beta pathology"

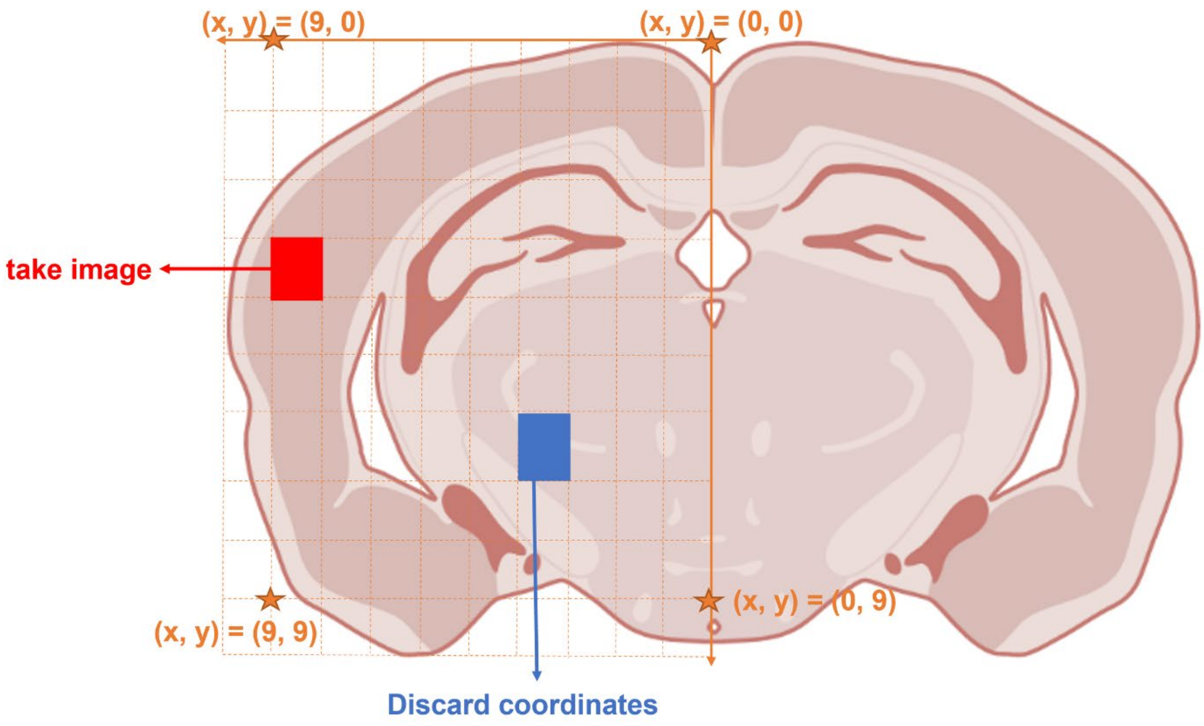


**S1 Fig. Schematic graph for the manual unbiased random selection method for confocal image acquisition.** This schematic graph shows the logic of picking coordinates for confocal images acquisition. The right hemisphere of the brain was equally divided into ten parts in both x and y directions. (x, y) coordinates were randomly generated using a random number generator. The microscope stage was moved to the target (x, y) coordinates using the stage rulings. When (x, y) coordinates fell into the areas of cortex, images were taken and cropped to include only cortical areas. When (x, y) coordinates fell out of the cortex the coordinates were discarded and no images were taken. Source of the mouse brain: biorender.com.

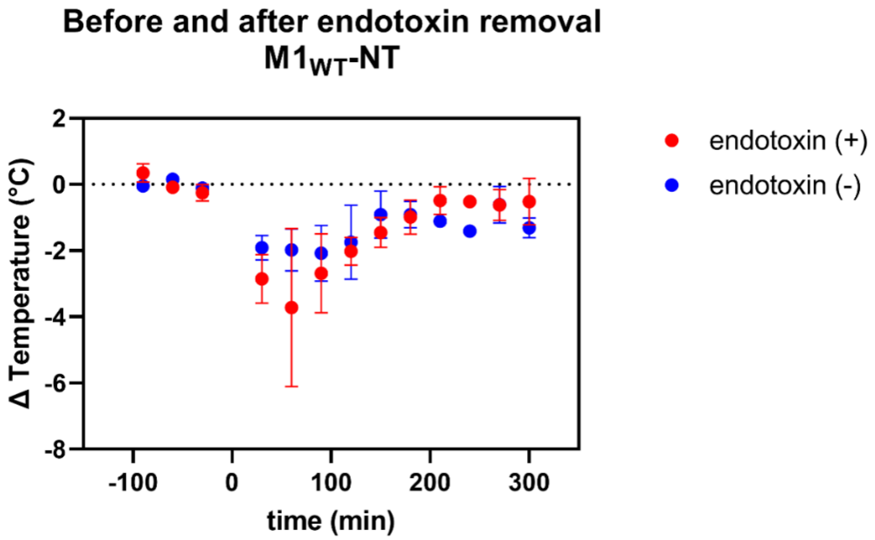


**S2 Fig. The hypothermia effect of M1WT-NT before and after endotoxin removal.** Endotoxin removal from in M1WT-NT preparations from E. coli decreased hypothermia effects at time 30min, 1hr and 1.5hr after IV injection of the M1 WT -NT (n= 3 per group). The maximum hypothermic effect of 600nmol/kg body weight of M1 WT -NT was approximately 2C° less after endotoxin removal.

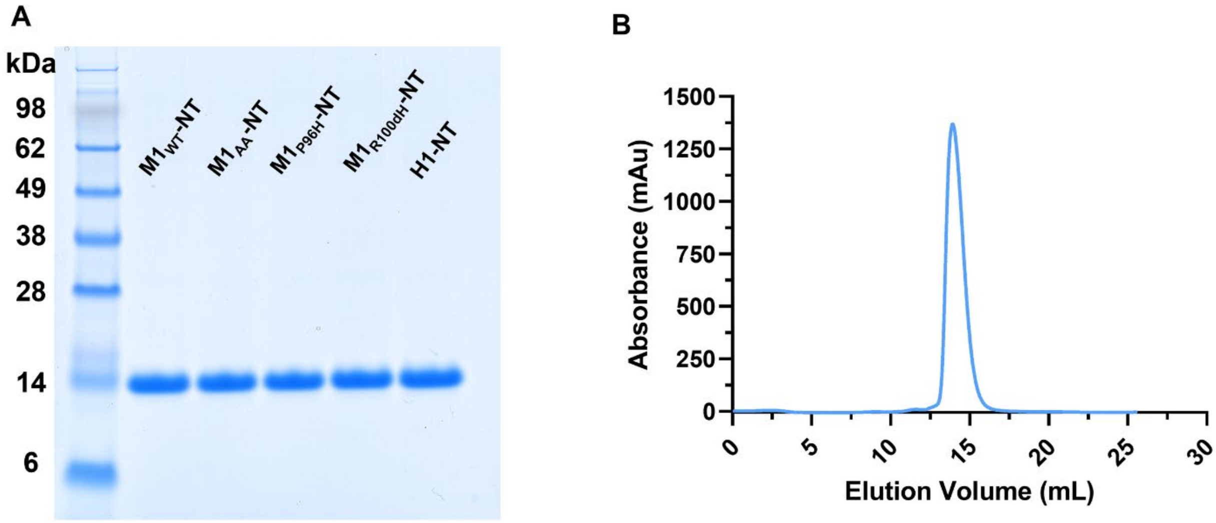


**S3 Fig. Post purification assessment of the five VHH-NT fusions M1WT-NT, M1P96H-NT, M1AA-NT, M1R100dH-NT and H1-NT.** The purity of the monomer VHH by (a) SDS-PAGE gel and exemplar (b) size-exclusion chromatography over a Superdex75 column for M1WT-NT indicating >95% purity and homogeneity of the material.

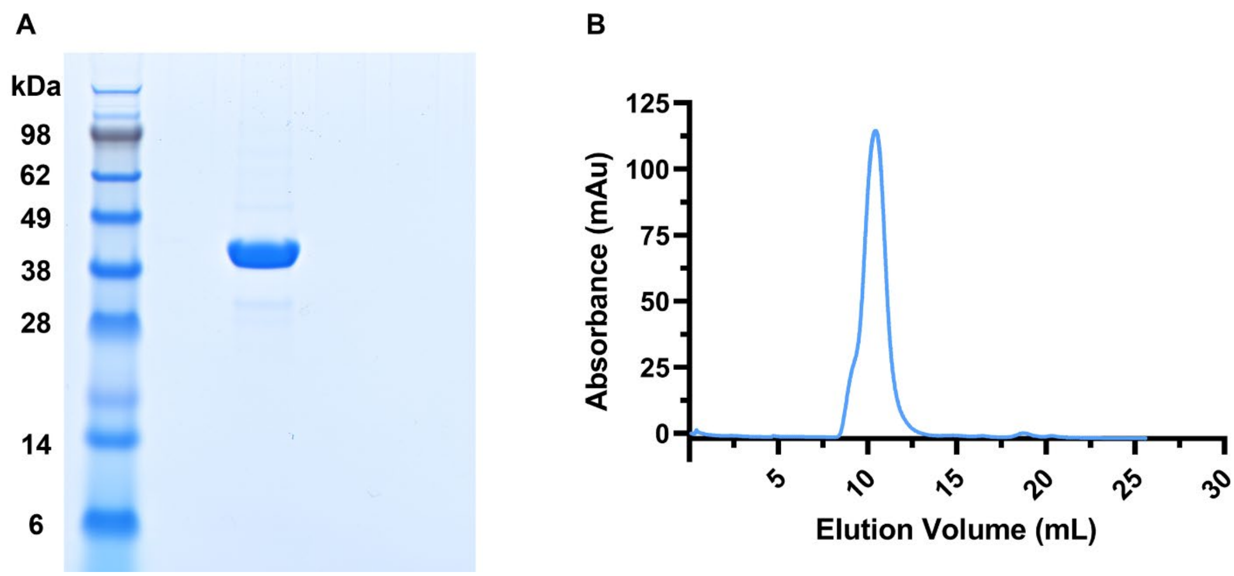


**S4 Fig. Post purification assessment of M1P96H-triplet-NT.** The purity of the triplet VHH by (a.) SDS-PAGE gel and exemplar (b.) size-exclusion chromatography over a Superdex75 column for M1P96H-Triplet-NT indicating >95% purity and homogeneity of the material.

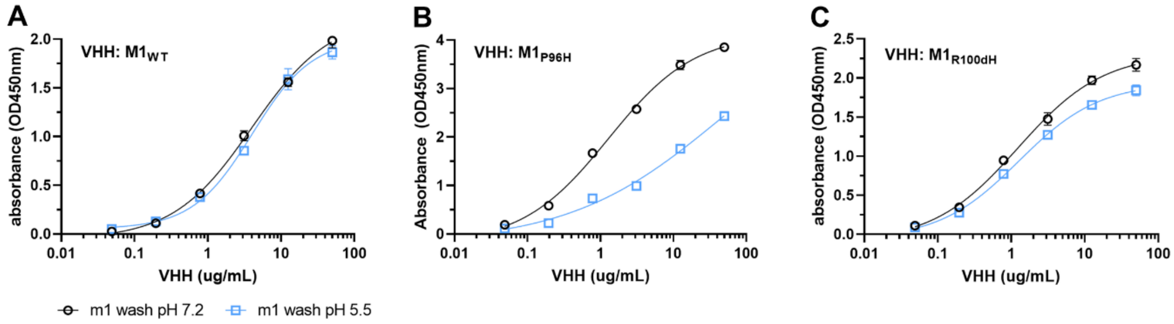


**S5 Fig. Histidine mutations impart a pH dependent dissociation effect in M1 VHH variants.** M1 variants (a.) M1WT, (b.) M1P96H and (c.) M1R100dH were incubated on mTfR coated ELISA plates, followed by a stringent wash with 1x PBS buffer at pH 7.2 or pH 5.5. Following pH dependent washing, the bound VHH was detected with an anti-alpaca-peroxidase antibody and the reaction terminated by addition of 1M HCl. Error bars represent the standard deviation of the mean values at each data point.

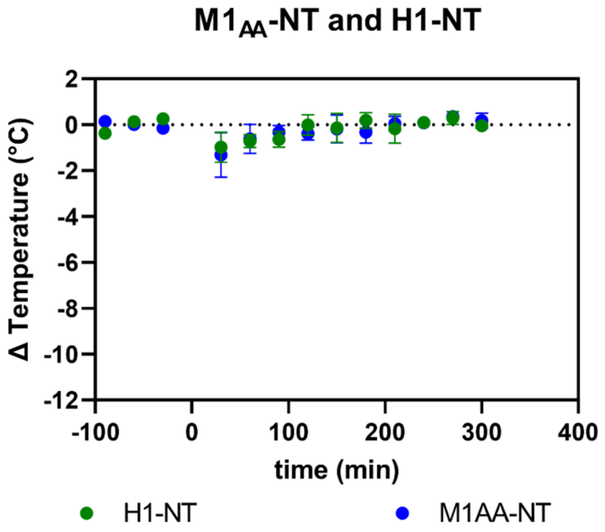


**S6 Fig. Lack of hypothermic effect with mTfR non-binding VHH-NT fusions.** M1AA-NT and H1-NT were injected at doses of 1400nmol/kg body weight. M1AA-NT and H1-NT lack prominent hypothermia effects (n=3 per group).

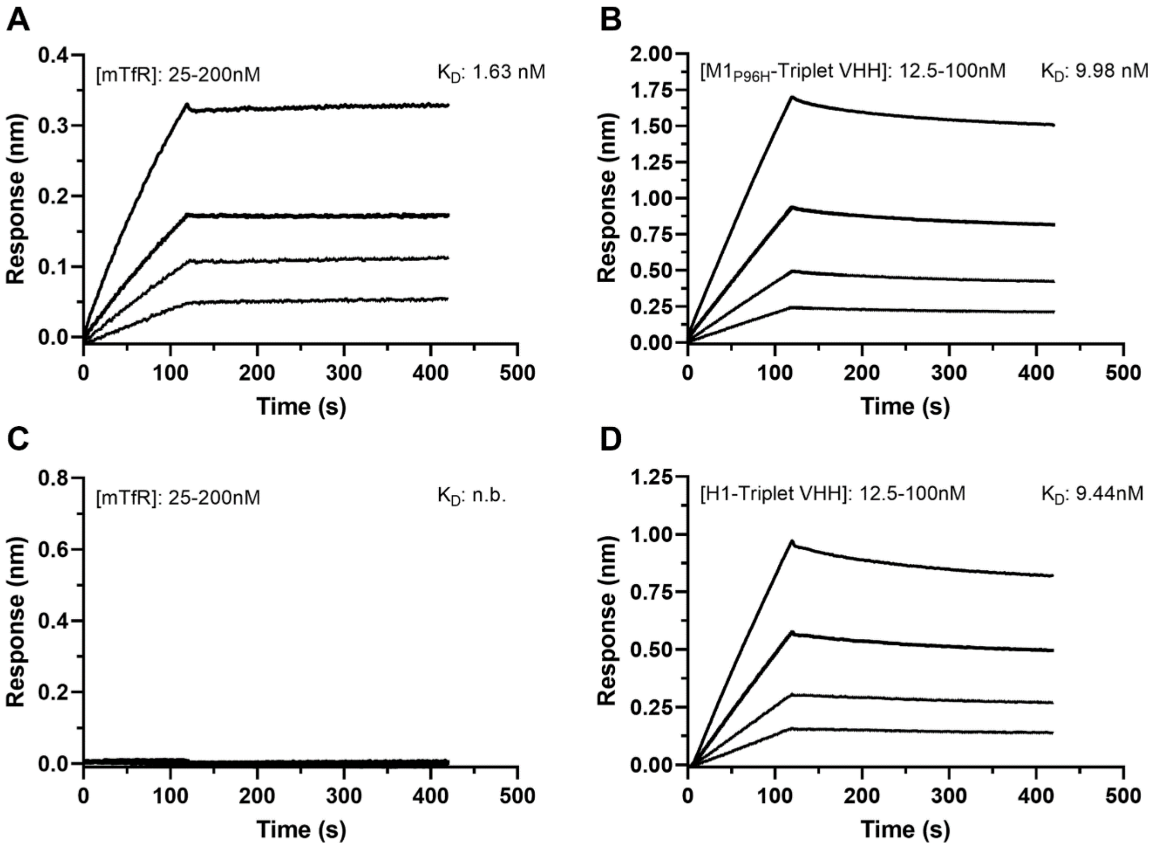


**S7 Fig. Affinity binding curves of M1P96H-triplet-NT and H1-triplet-NT binding affinity (KD) to mTfR and Aβ.** Using biolayer interferometry on an Octet Red96 system, association and dissociation rates were determined by immobilizing biotinylated VHH (a, c) or biotinylated amyloid beta (b, d) onto streptavidin-coated optical sensors. M1P96H-triplet-NT association and dissociation curve to (a) mTfR and (b) Aβ. The affinity of M1P96H-triplet-NTwas measured to be 1.63nM to mTfR and 9.98nM to Aβ. H1-triplet-NT association and dissociation curve to (c) mTfR and (d) Aβ. The affinity of H1-triplet-NT was measured to be 9.44nM to Aβ and no binding to mTfR. (n.b. = no binding).

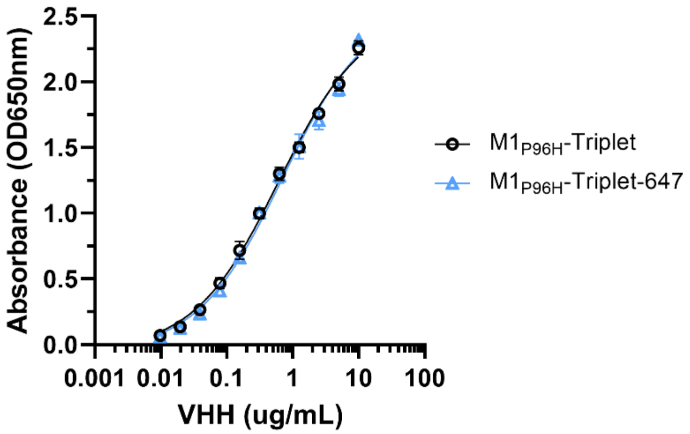


**S8 Fig. mTfR ELISA on M1P96H-Triplet before and after 647 labelling.** VHH was incubated with mTfR absorbed to ELISA plates and detected using an anti-alpaca-peroxidase antibody to determine the impact of fluorophore conjugation. The near overlapping binding curves indicate a lack of effect following fluorophore labelling. Error bars represent the standard deviation of the mean values at each data point.

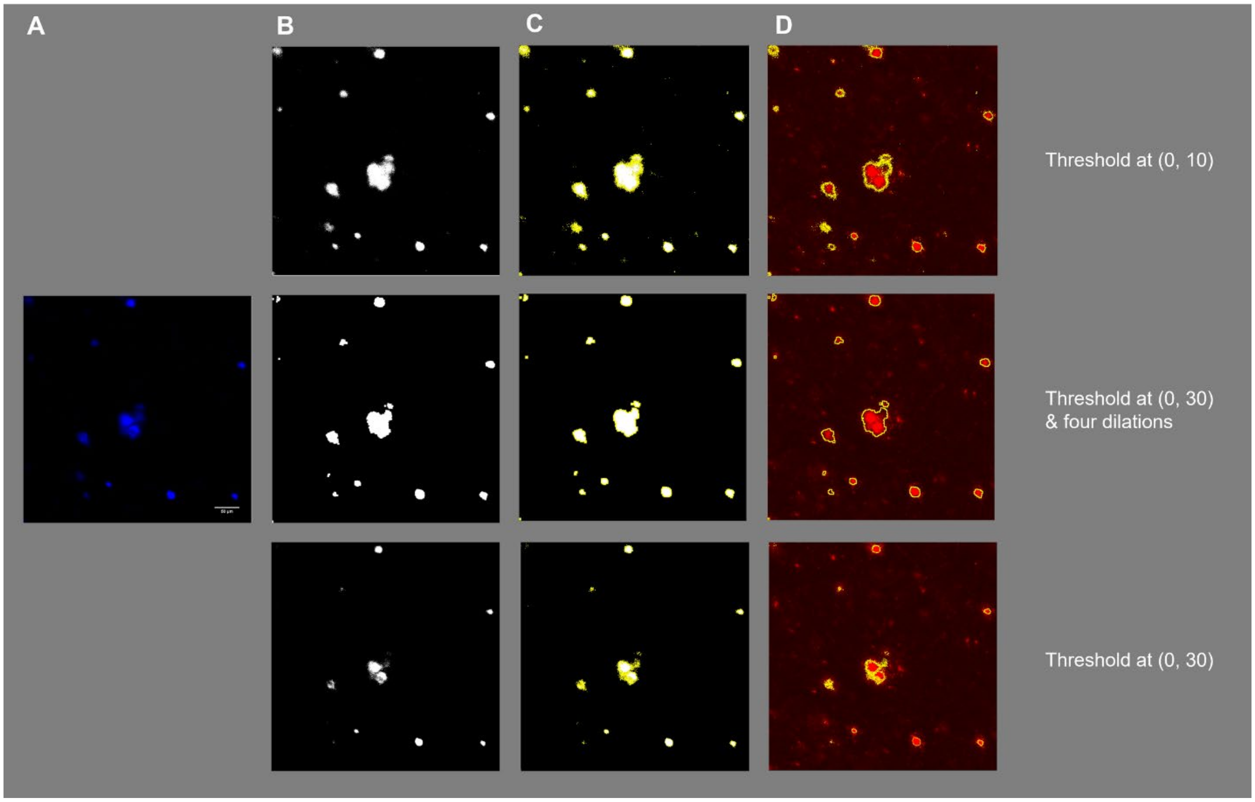


**S9 Fig. Quantitative analysis procedure for confocal images with thresholding followed by automated plaque selection and mean 647 channel signal calculation.** An APP/PS1 positive naïve brain slice with ex vivo M1P96H-Triplet-647 labeling was used as the example. Column a. shows a representative confocal microscope image of cortex X34 staining. Column b. shows the X34 labeled images after thresholding.

Column c. shows automated selection of regions of plaques based on the thresholded images. Column d. shows the application of selected areas (from thresholded X34 images) to the 647 channel images.
